# Development of quantitative high-throughput screening assays to identify, validate, and optimize small-molecule stabilizers of misfolded β-glucocerebrosidase with therapeutic potential for Gaucher disease and Parkinson’s disease

**DOI:** 10.1101/2024.03.22.586364

**Authors:** Darian Williams, Logan M. Glasstetter, Tiffany T. Jong, Abhijeet Kapoor, Sha Zhu, Yanping Zhu, Alexandra Gehrlein, David J. Vocadlo, Ravi Jagasia, Juan J. Marugan, Ellen Sidransky, Mark J. Henderson, Yu Chen

## Abstract

Glucocerebrosidase (GCase) is implicated in both a rare, monogenic disorder (Gaucher disease, GD) and a common, multifactorial condition (Parkinson’s disease); hence, it is an urgent therapeutic target. To identify correctors of severe protein misfolding and trafficking obstruction manifested by the pathogenic L444P-variant of GCase, we developed a suite of quantitative, high-throughput, cell-based assays. First, we labeled GCase with a small pro-luminescent HiBiT peptide reporter tag, enabling quantitation of protein stabilization in cells while faithfully maintaining target biology. TALEN-based gene editing allowed for stable integration of a single HiBiT-*GBA1* transgene into an intragenic safe-harbor locus in *GBA1*-knockout H4 (neuroglioma) cells. This GD cell model was amenable to lead discovery via titration-based quantitative high-throughput screening and lead optimization via structure-activity relationships. A primary screen of 10,779 compounds from the NCATS bioactive collections identified 140 stabilizers of HiBiT-GCase-L444P, including both pharmacological chaperones (ambroxol and non-inhibitory chaperone NCGC326) and proteostasis regulators (panobinostat, trans-ISRIB, and pladienolide B). Two complementary high-content imaging-based assays were deployed to triage hits: the fluorescence-quenched substrate LysoFix-GBA captured functional lysosomal GCase activity, while an immunofluorescence assay featuring antibody hGCase-1/23 provided direct visualization of GCase lysosomal translocation. NCGC326 was active in both secondary assays and completely reversed pathological glucosylsphingosine accumulation. Finally, we tested the concept of combination therapy, by demonstrating synergistic actions of NCGC326 with proteostasis regulators in enhancing GCase-L444P levels. Looking forward, these physiologically-relevant assays can facilitate the identification, pharmacological validation, and medicinal chemistry optimization of new chemical matter targeting GCase, ultimately leading to a viable therapeutic for two protein-misfolding diseases.

**Significance Statement:** Gaucher disease, the inherited deficiency of glucocerebrosidase, is caused by biallelic, loss-of-function mutations in the gene *GBA1,* which is also the most frequent genetic risk factor for Parkinson’s disease. While the development of small-molecule stabilizers of glucocerebrosidase is being considered for both disorders, discovery and optimization of lead compounds is limited by the lack of robust cell-based assays amenable to high-throughput screening format. We developed a comprehensive assay pipeline for preclinical discovery of glucocerebrosidase modulators and began by screening libraries enriched with bioactive compounds with known mechanisms of action. The screen identified chemical matter with established relevance to glucocerebrosidase, provided an atlas of potential new molecular targets regulating the *GBA1* pathway, and produced a set of promising potential therapeutics.

## Introduction

*GBA1* encodes β-glucocerebrosidase (GCase), a lysosomal enzyme that hydrolyzes glycosphingolipid substrates, including glucosylceramide (GluCer) and glucosylsphingosine (GluSph). Biallelic loss-of-function mutations in *GBA1* cause Gaucher disease (GD), the most common lysosomal storage disorder (LSD). In addition, *GBA1* mutations serve as the most frequent genetic risk factor for synucleinopathies, such as Parkinson’s disease (PD) [1] and dementia with Lewy bodies (DLB) [2]. This genetic association is strongly supported by preclinical [3, 4] and clinical [5, 6] evidence. GCase is thus firmly positioned as a well-validated therapeutic target for modification of both a rare, monogenic disorder and a common, multifactorial disease, motivating translational efforts aimed at its enhancement.

The current standard-of-care for GD is enzyme replacement therapy (ERT), whereby recombinant, mannose-terminated GCase is delivered to lipid-laden macrophages via biweekly intravenous infusions. While ERT offers dramatic reversal of the peripheral symptoms of the disease, such as hepatosplenomegaly and hematologic abnormalities [7], it is rapidly cleared from the blood and, unfortunately, does not penetrate the blood-brain barrier (BBB) [8]. Therefore, it has no effect on the neurologic manifestations seen in type 2 (acute neuronopathic) or type 3 (subacute neuronopathic) GD [9], and it does not prevent or modify the progression of parkinsonism in patients with GD [10]. In addition, ERT is prohibitively expensive (with an average lifetime cost of over $6,000,000 per patient [11]) and inconvenient. Since GD is a multi-systemic disorder, an oral small-molecule therapy would be highly desirable. Unfortunately, substrate reduction therapy, which blocks the formation of pathological substrates via GluCer synthase inhibitors, does not alter the progression of *GBA1*-PD [12], and it fails to target GCase dysfunction upstream of the lipid pathology. Thus, there is an urgent unmet need for a BBB-penetrant, orally available small-molecule drug that enhances GCase levels and functions in the brain. We envision that such a pharmaceutical would be widely applicable, as a first-line therapy for neuronopathic GD, as a prospective prophylactic agent against *GBA1*-PD, and, perhaps most broadly impactful, as a potential disease-modifying therapy for sporadic PD [3, 13–15].

In most cases, GD is fundamentally a loss-of-function protein-misfolding disease, like cystic fibrosis, Fabry disease, and GM1 gangliosidosis [16]; the underlying mechanism is not aggregation but premature endoplasmic reticulum-associated degradation (ERAD) [17–19] of misfolded, missense-mutant GCase, which retains some residual catalytic activity if properly trafficked to the lysosome [20–22]. The ERAD process involves recognition and retention of aberrant GCase in the ER, followed by its retro-translocation into the cytosol and subsequent degradation by the ubiquitin-proteasome system [23]. Importantly, different mutant GCase variants present variable degrees of ERAD, underlying some of the clinical heterogeneity encountered in GD [24]. The two most common *GBA1* mutations are N370S (p.N409S), which represents 70% of the mutant alleles in Ashkenazi Jews and is exclusively associated with type 1 GD, and L444P (p.L483P), a severe, pan-ethnic mutation on the hydrophobic core of the Ig-like domain of GCase that leads to substantial protein instability and is often encountered in neuronopathic (types 2 and 3) GD [25]. Severely-misfolded GCase variants like L444P consume valuable proteostasis network capacity within the cell, contributing to a potential gain-of-toxic-function phenotype that could link to PD [26].

Attempts to salvage misfolded GCase from the ERAD pathway using small molecules – in the form of enzyme enhancement therapy (EET) – have focused on two potentially synergistic therapeutic paradigms [27]: a generic biological arm involving reprogramming of the protein homeostasis network with proteostasis regulators (PRs) [28–32], and a tailored chemical arm consisting of pharmacological chaperones (PCs) that thermodynamically stabilize the native state of the enzyme [33] or mobilize active dimer formation [34–36] through direct binding. The latter approach initially focused on the identification of active-site inhibitors; these are primarily substrate-mimetic compounds (iminosugars), such as isofagomine [37, 38] and *N*-(*n*-nonyl)deoxynojirimycin [33], but some unique chemical scaffolds were also identified via high-throughput screening efforts [39–41]. These compounds, which must be utilized at sub-inhibitory concentration, suffer from a narrow therapeutic window between chaperoning and inhibitory behavior, and they exhibit poor selectivity against related hydrolases (e.g., α-glucosidase, α-galactosidase) [42]. The most promising PC in the inhibitor class is ambroxol, an expectorant that was isolated in a screen of 1,040 approved drugs based on a thermal denaturation assay with wild-type recombinant GCase [43]. Ambroxol exhibits pH-dependent chaperone behavior [43], is BBB-permeable and well-tolerated [13, 44, 45], and showed favorable results in small pilot studies for neuronopathic GD [45, 46], with clinical trials in progress for PD [47] and DLB [48].

The (pre-)clinical promise of ambroxol notwithstanding, it is an inhibitory chaperone [49], which places a ceiling on its therapeutic potential [13]. Non-inhibitory, allosteric-site directed PCs have emerged as a favorable therapeutic strategy, with a wider therapeutic window than active-site inhibitors. The first members of this class were discovered through a quantitative high-throughput screening (qHTS) campaign [50], which utilized Gaucher spleen homogenate as a source of mutant (N370S) GCase [51] and fluorogenic substrate to read out activity. Two hits from this screen of 250,000 compounds were advanced by medicinal chemistry to yield lead compounds NCGC607 [52] and NCGC758 [53, 54], which have distinct chemotypes. In *GBA1*-PD patient-derived macrophages, these PCs increased GCase protein levels, enhanced GCase activity, and reduced GluCer and GluSph substrate accumulation; in patient-derived dopaminergic neurons, they also lowered α-synuclein levels [52, 53, 55].

While the aforementioned *in vitro* qHTS assay [51] enables biochemical evaluation of PCs that compose the chemical arm of EET, it is not useful for identifying small-molecule PRs to advance the biological arm. Furthermore, the binding of small molecules to GCase, modulation of its enzymatic activity, and promotion of its folding do not always correlate, so previously-described screening methods are not adequate to drive medicinal chemistry efforts toward optimization of PCs. Ideally, lead compounds should be discovered and optimized in a cellular or phenotypically-relevant context. In this work, we solve these issues by successfully labeling GCase with a small (1.3 kDa) HiBiT tag [56], which undergoes high-affinity complementation with exogenous LgBiT to reconstitute an active luciferase. As a primary assay, HiBiT provides a bioluminescent readout of GCase protein levels in cells that is amenable to both lead discovery via qHTS [57–59] and medicinal chemistry optimization of leads via structure-activity relationships. We elected to integrate this targeted phenotypic approach into a drug screening funnel [60]. To confirm that hits identified from the primary screen enhance GCase lysosomal activity and trafficking, we implemented two high-content imaging-based secondary assays, leveraging the fluorescence-quenched substrate LysoFix-GBA [61] and the newly-described GCase antibody hGCase-1/23 [62]. Finally, as an endpoint, we evaluated the reversal of glycosphingolipid substrate accumulation [63]. We chose to first deploy this screening pipeline on pharmacologically-active, mechanistically-annotated compound libraries [64, 65], with a focus on drug repurposing [58] for the biological arm of EET [30]. The best PRs derived from this screen were then combined with a newly-identified analog of an in-house, non-inhibitory PC, to establish a potent and efficacious synergistic co-formulation [27]. Future work will deploy the pipeline on a diversity library, in order to discover novel PCs and PRs.

## Results

### The HiBiT peptide reporter tag preserves trafficking and function of labeled GCase variants, providing a cell-based platform for qHTS

The HiBiT reporter is an 11-amino acid peptide tag suitable for luminescence-based qHTS. To explore the feasibility of tagging human GCase at its N-terminus with HiBiT (**Fig. 1A**), prior to generating stable cell lines, we transiently transfected cells with constructs encoding a transgene containing the *GBA1* signaling peptide, the HiBiT peptide tag, and a Gly/Ser linker upstream of *GBA1*. The transgene was expressed in a *GBA1*-KO H4 (human neuroglioma) cell line, and localization of the reporter protein was assessed using a monoclonal human GCase antibody, hGCase-1/23 [62] (**Fig. S1**). When moderately expressed, HiBiT-GCase-WT showed specific lysosomal localization comparable to endogenous GCase and transfected untagged GCase, suggesting that the HiBiT tag did not interfere with GCase trafficking. However, higher levels of untagged GCase led to ER accumulation without clear lysosomal localization, indicating that GCase trafficking is perturbed by overexpression. To address this, HiBiT-GCase-WT expression was constrained by integrating the transgene cassette into a safe-harbor site within an intron of the Citrate Lyase Beta-Like (*CLYBL*) gene using transcription activator-like effector nucleases (TALENs) [66, 67] in the *GBA1*-KO H4 cell line (**Fig. 1B**). This integrative gene transfer method allows efficient knock-in of the large transgene, with minimal impact on local and global gene expression [66]. Sustained expression of the transgene is driven by a strong CAG promoter. A *GBA1*-KO H4 clone with a single copy of HiBiT-*GBA1*-WT was identified via Droplet Digital PCR (ddPCR) and used for subsequent experiments (**Fig. 1D**). The expression level (**Fig. 1E,G**) and activity (**Fig. 1H**) of HiBiT-GCase-WT were moderately higher than those of endogenous GCase. Both proteins showed similar lysosomal localization (**Fig. 1C**) and glycosylation status (**Fig. 1F**). Additionally, HiBiT-GCase-WT successfully reduced accumulated GluSph in the *GBA1*-KO H4 cell line (**Fig. 1I**), behaving akin to endogenous GCase and confirming functional activity of the reporter protein.

**Figure 1.**
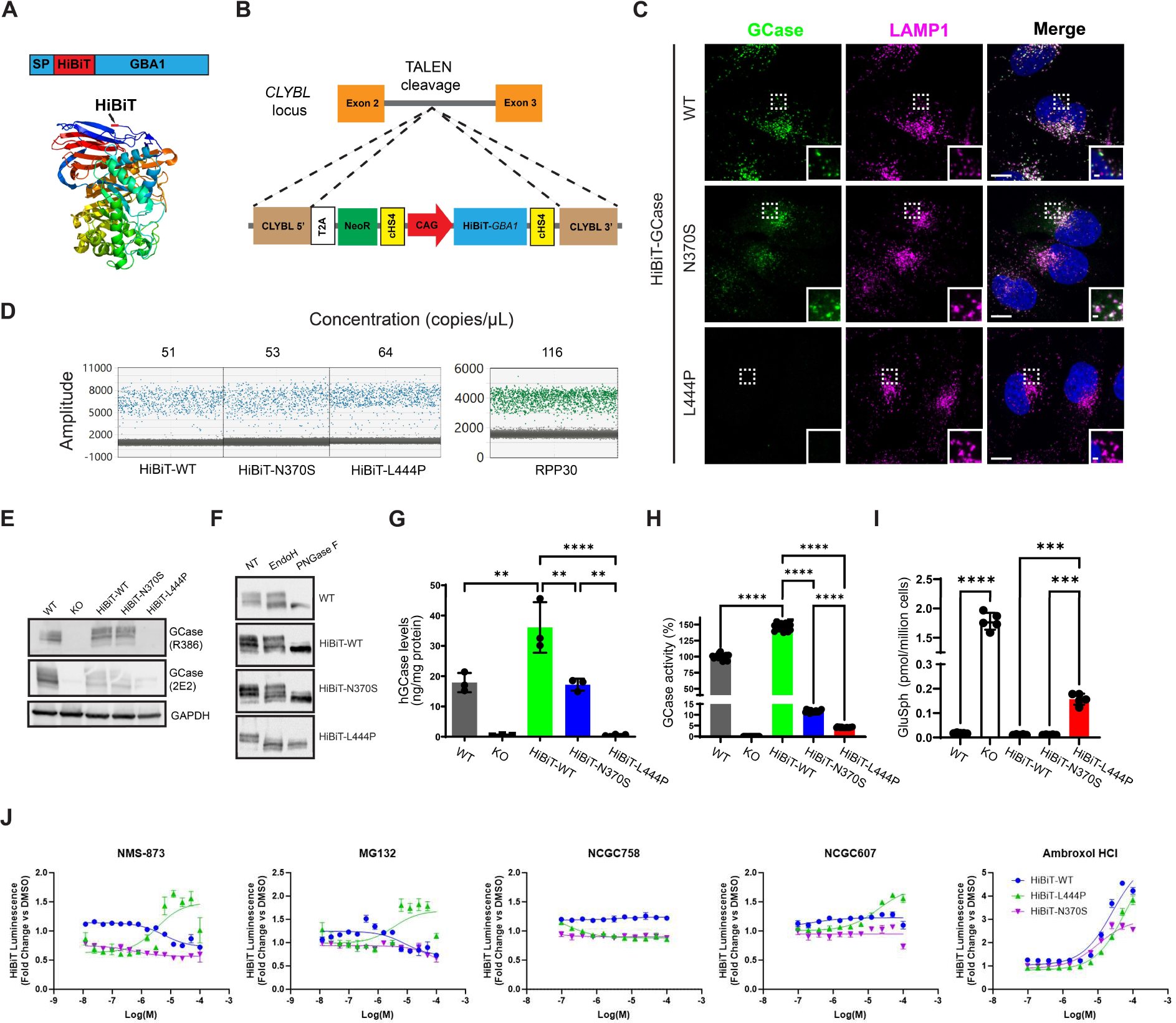
HiBiT-tagged GCase retains normal trafficking and function, enabling a high-throughput screening assay measuring cellular GCase levels. (**A**) GCase was labeled with a small (1.3 kDa), pro-luminescent, N-terminal HiBiT peptide tag immediately following the signal peptide (SP) sequence. (**B**) This HiBiT-GCase reporter (WT, N370S, or L444P) was engineered into a human Citrate Lyase Beta-Like (*CLYBL)* intragenic safe-harbor locus within a *GBA1*-KO H4 cell background using TALEN-enhanced integrative gene transfer. (**C**) Co-localization of GCase (green) with lysosomal marker LAMP1 (magenta) was determined by immunofluorescent staining. (**D**) The copy number of the stably-integrated transgene was confirmed to be 1 across all three HiBiT-GCase lines via Droplet Digital polymerase chain reaction (ddPCR). The HiBiT-GCase H4 lines featured ∼ 60 copies/μL of the HiBiT-*GBA1* transgene, as compared with ∼ 120 copies/μL of the reference gene RPP30, which has a known copy number of 2 in the *GBA1*-WT H4 cell line. (**E**) GCase protein level was measured by Western blot in *GBA1*-WT, *GBA1*-KO, and HiBiT-GCase H4 cell lines using anti-GCase (R386, 2E2) antibodies. (**F**) Glycosidase sensitivity analysis indicates that HiBiT-GCase-L444P is entirely retained in the ER. The Endo H-sensitive fraction (lower band) on the blot contains immature, ER-retained GCase, while the Endo H-resistant fraction (top band) contains maturely-glycosylated, post-ER-localized GCase. Both fractions are responsive to PNGase F treatment. NT: non-treated. (**G**) GCase protein levels were quantitated by AlphaLISA (Amplified Luminescent Proximity Homogeneous Assay) utilizing a sandwich configuration of two monoclonal antibodies recognizing non-overlapping epitopes, hGCase-1/23 (which was biotinylated and associated with a streptavidin-coated donor bead) and hGCase-1/17 (which was directly conjugated to an acceptor bead). (Error bars: SEM [*n* = 3 biological replicates]). (**H**) GCase activity was measured in cell lysates using the fluorogenic substrate 4-methylumbelliferyl-β-D-glucopyranoside. Relative GCase activity was calculated by adjusting for protein concentration, correcting for *GBA1*-KO H4 cell background, and normalizing to *GBA1*-WT signal. (Error bars: SD [*n* = 16 technical replicates]). (**I**) Levels of glucosylsphingosine (GluSph) in H4 cell pellets were quantified by positive ion electrospray LC-MS/MS in multiple reaction-monitoring mode, using deuterated compounds as internal standards. (Error bars: SD [*n* = 5 biological replicates]). (**J**) Pilot testing of the HiBiT-GCase assay was performed in ultra-high-throughput 1536-well plate format. Cells were treated with known-active ERAD modulators (NMS-873, p97 inhibitor; MG132, proteasome inhibitor) or GCase stabilizers (ambroxol, NCGC758, or NCGC607) for 24 h, followed by measurement of HiBiT-GCase luminescence. For each respective cell line, data are represented as fold change in luminescence (RLU) in compound-treated versus DMSO-treated cells. (Error bars: SEM [*n* = 3 – 6]). Dose-response curves were fit using log(agonist) vs. response (three parameters). *P-value ≤ 0.05; **P-value ≤ 0.01; ***P-value ≤ 0.001; ****P-value ≤ 0.0001.

Following this success, HiBiT-GCase-N370S and -L444P lines were generated using the same strategy in *GBA1*-KO H4 cells. HiBiT-GCase-N370S exhibited lysosomal localization similar to HiBiT-GCase-WT, whereas HiBiT-GCase-L444P had no staining, likely due to severe misfolding (**Fig. 1C**). The expression level of HiBiT-GCase-L444P, detected by Western blot, was notably lower than HiBiT-GCase-WT and HiBiT-GCase-N370S levels (**Fig. 1E**). GCase protein levels in the HiBiT-GCase-N370S and -L444P lines were quantified by a bead-based immunoassay – Amplified Luminescent Proximity Homogeneous Assay (AlphaLISA) – as 48% and 2% of HiBiT-GCase-WT levels, respectively (**Fig. 1G**). Endo H treatment revealed sensitivity in a small fraction of HiBiT-GCase-N370S and a large fraction of HiBiT-GCase-L444P, indicating the immature glycosylation status of the latter, resulting from increased misfolding and ER retention (**Fig. 1F**). GCase activity was reduced for both HiBiT-GCase-N370S and -L444P (**Fig. 1H**). While both HiBiT-GCase-WT and -N370S successfully eliminated accumulated GluSph in the *GBA1*-KO H4 cell line, HiBiT-GCase-L444P only partially decreased the accumulation (**Fig. 1I**).

### HiBiT-GCase-L444P is responsive to PCs and PRs in a quantitative high-throughput luminescence assay

To demonstrate the utility of the HiBiT-GCase H4 lines as a drug discovery tool, we miniaturized the HiBiT assay to ultra-high-throughput (1536-well plate) format and tested a set of known-active GCase stabilizers after 24 h treatment. MG-132 is a proteasome inhibitor that also upregulates ER folding capacity through unfolded protein response (UPR) activation [27]. NMS-873 is an inhibitor of the Valosin-containing protein (VCP) p97, which hydrolyzes ATP to extract misfolded proteins from the ER into the cytosol, thereby enabling ERAD via the proteasome [68]. Both PRs selectively stabilized HiBiT-GCase-L444P with a dose-dependent response, indicating its constitutive loss through ERAD [24] (**Fig. 1J**). The response to non-inhibitory, allosteric-site-directed PC NCGC607 [52] displayed a similar selectivity for HiBiT-GCase-L444P, whereas the inhibitory, active-site-directed PC ambroxol [13, 43, 45, 69] increased HiBiT-GCase levels in all three lines. Notably, non-inhibitory PC NCGC758 was inactive in all three lines under these experimental conditions. Overall, these results demonstrate the ability of the HiBiT-GCase method to detect relevant small-molecule stabilizers of GCase, including both PRs and PCs. A treatment period of 24 h was sufficient to detect changes in the steady-state levels of GCase; biogenesis and maturation of native GCase occur within this timeframe [70], and the half-lives of N370S- and L444P-mutant GCase are only ∼6 h and ∼1 h, respectively [71]. While a fraction of GCase folding intermediates will be targeted for ERAD regardless of mutation status, ERAD plays a more direct, rate-limiting role in the processing of the severely-misfolded L444P variant, as compared with WT or N370S [72]. Under the given assay conditions, the effect of PRs on GCase stabilization was specific to the L444P reporter line, so this neuronopathic variant was the focus of subsequent high-throughput screening.

### Identification of novel compounds and mechanistic classes that promote stabilization of HiBiT-GCase-L444P protein levels

Using the HiBiT-GCase-L444P H4 post-translational reporter cell line (**Fig. 2A**), we performed qHTS on a collection of 10,779 compounds across three annotated small-molecule libraries: NPC (NCATS Pharmaceutical Collection) [64], NPACT (NCATS Pharmacologically Active Chemical Toolbox) [65], and HEAL (Helping to End Addiction Long-term; https://ncats.nih.gov/research/research-activities/heal/expertise/library). We also screened analogs of two non-inhibitory PC chemotypes, NCGC607 and NCGC758, based on a SMILES similarity search within all internal NCATS libraries using a similarity cutoff of 80%, which retrieved 187 compounds (NCGC607 analogs: 91 compounds; NCGC758 analogs: 96 compounds). All 10,779 compounds were screened at 7 concentrations ranging most commonly from 5 nM – 75 µM (**Fig. 2B**). From the primary screen, 716 compounds (6.6%) met the selection criteria as hits and were cherrypicked for expanded dose-response testing, along with CellTiter-Glo cytotoxicity testing (**Fig. S2**). Of the cherrypicked hits, 140 were confirmed in follow-up testing: 75 final hits were derived from NPACT, 43 from HEAL, 6 from NPC, and 16 from the set of chaperone analogs (**Fig. 2B; Table S1**). The final hits represented a few salient mechanisms of action (**Fig. 2C**), and analysis of their molecular targets (**Fig. S3**) revealed several enriched pathways (**Table S2**). Most hits were epigenetic modulators, including histone deacetylase (HDAC) inhibitors (42/140) like panobinostat, and BET bromodomain protein inhibitors or degradation inducers (5/140) like ARV-825. HDAC inhibitors are known GCase-L444P stabilizers [73, 74]. In addition, known actives NMS-873 (**Fig. S4**) and ambroxol HCl (**Fig. 2C**) were detected as final hits in the screen, providing strong prospective validation of the HiBiT-GCase-L444P H4 reporter cell line as a powerful qHTS tool that identifies relevant chemical matter.

**Figure 2.**
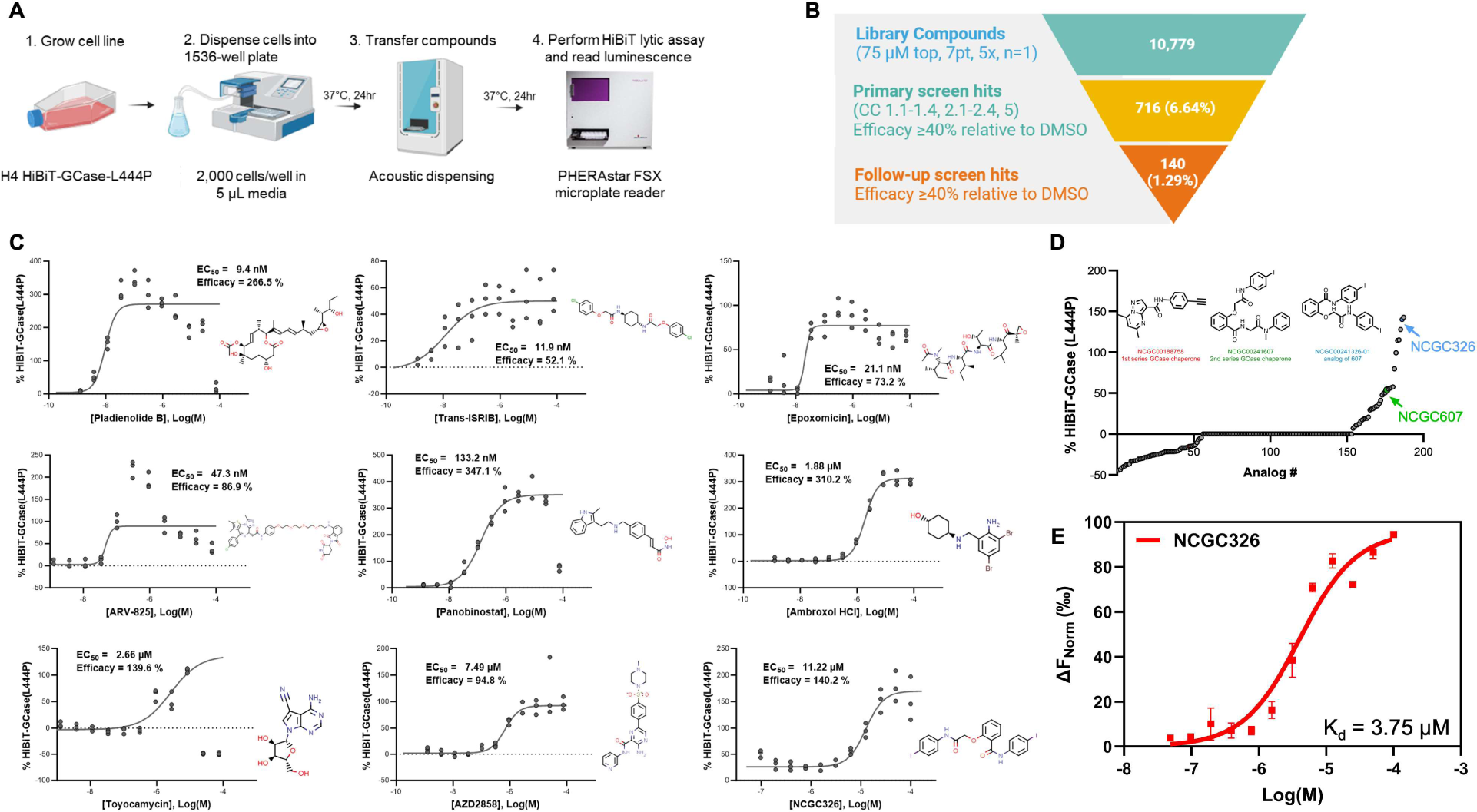
Quantitative high-throughput screening of mechanistically-annotated small-molecule libraries for GCase-L444P stabilizers. (**A**) Schematic of high-throughput screening methodology using H4 HiBiT-GCase-L444P reporter cell line. (**B**) In the primary screen, 10,779 compounds, including the NCATS Pharmaceutical Collection (NPC), NCATS Pharmacologically Active Chemical Toolbox (NPACT), and Helping to End Addiction Long-term (HEAL) chemical libraries, as well as analogs of the non-inhibitory chaperone chemotypes NCGC607 and NCGC758, were evaluated at concentrations most commonly ranging from 5 nM – 75 μM in a 7-point, 5x dilution series using a single replicate. Compounds were then triaged based on curve class (CC) and efficacy: those with CC of 1.1 – 1.4, 2.1 – 2.4, or 5 and efficacy ≥ 40% were considered primary screen hits. These 716 compounds were retested (NCGC326: 100 nM – 100 μM, 11-point, 2x dilution; others: 1 nM – 75 μM, 11-point, 3x dilution) with *n* = 3 replicates and categorized as final hits if they met a cutoff of efficacy ≥ 40%, regardless of curve class, resulting in 140 confirmed hits. (**C**) Top representative follow-up screen hits from each mechanistic cluster were selected based on their potency and efficacy in stabilizing GCase-L444P levels. Response values were normalized to intraplate DMSO-treated controls, such that 100% efficacy reflects a doubling of GCase levels. *n* = 3. (**D**) Dose-response profiles for 187 analogs of NCGC758 and NCGC607 revealed 11 compounds, including NCGC326 (blue), with greater efficacy than NCGC607 (green). Chaperone NCGC758 was inactive under the screening conditions. (**E**) Target engagement studies with microscale thermophoresis (MST) revealed that chaperone NCGC326, a follow-up screen hit and analog of NCGC607, binds recombinant GCase-WT with a dissociation constant (K_d_) of 3.75 μM in 50 mM sodium citrate buffer (pH 5.5). *n* = 2.

Other mechanistic classes identified in the primary screen included two proteasome inhibitors (epoxomicin and ixazomib) and five glycogen synthase kinase 3 (GSK-3) inhibitors [75] like AZD2858 (**Fig. 2C**). Of note, the most potent hit identified in the primary screen was pladienolide B (EC_50_ = 9.4 nM), a splicing factor SF3B1 modulator (**Fig. 2C**). The third-most potent hit was trans-ISRIB (**Fig. 2C**), a known PR acting through inhibition of the PERK-mediated UPR pathway, whereby it releases the brake on translation [76]; its potential therapeutic utility for GD or *GBA1*-PD has not been previously investigated. Toyocamycin is another hit acting through UPR inhibition; specifically, it blocks activation of the IRE1α-XBP1 arm [77]. The most efficacious hit was an HDAC inhibitor, vorinostat (SAHA), with an efficacy of 658% relative to the DMSO control (**Table S1**; **Fig. S4**). The screen of analogs of PCs NCGC607 and NCGC758 (**Fig. 2D**) captured thirteen active compounds related to the NCGC607 (salicylic acid derivative) scaffold, including NCGC607 itself (**Fig. S4**), and three active compounds with structural similarity to the NCGC758 (pyrazolopyrimidine) scaffold, including the investigational drug LTI-291 / BIA 28-6156 (**Fig. S4**), which is currently being evaluated in a clinical trial for *GBA1*-PD [78, 79]. None of the PC analogs showed improved potency relative to the active parent compound NCGC607 (EC_50_ > 10 μM for all compounds). Nonetheless, the screen identified NCGC326 as a novel analog of NCGC607 with ∼3-fold increased efficacy (140% vs. 54%) and similar potency (**Fig. 2C**). Moreover, microscale thermophoresis (MST) [54] validated the binding affinity of NCGC326 for recombinant GCase-WT (K_d_ = 3.75 μM), reaffirming its status as a PC (**Fig. 2E**). Finally, selected final hits underwent a counter-screen to rule out interference with the reconstituted luciferase enzyme during the HiBiT-GCase-L444P assay in H4 cells (**Fig. S5**). Representative hits from each mechanistic cluster, along with the newly-discovered PC NCGC326, were subsequently analyzed in orthogonal secondary assays.

### Orthogonal high-content screening assays enable triage of hits from the primary screen

To enable triage of hits from the primary screen, we next sought to develop high-content imaging-based secondary assays to characterize the effect of small molecules on lysosomal translocation of an enzymatically active GCase. Small-molecule regulators of GCase-L444P can increase cellular protein levels without enhancing lysosomal translocation and enzymatic activity of the mutant protein. For example, proteasome inhibitors like bortezomib may not relinquish persistent ER-retention of the misfolded protein, while active-site-directed PCs like isofagomine (and even ambroxol) have an inhibitory effect on GCase lysosomal activity at high concentrations (μM) [49]. Therefore, compounds which can increase total levels of GCase-L444P while simultaneously increasing lysosomal translocation and functional activity offer the greatest therapeutic potential. To identify such compounds, we implemented two complementary high-content screening (HCS) assays [60, 80–82]. First, a *GBA1*-specific, fluorescence-quenched substrate, LysoFix-GBA, was used to directly evaluate GCase function within lysosomes [61]. This sensitive lysosomal activity probe is selective against immature GCase forms in the ER and Golgi apparatus, as well as against other β-glucosidases [61]. It is also fixable, lysosomotropic (to minimize diffusive signal loss), and red-shifted (to reduce autofluorescence background) (**Fig. S6A**) [61]. Second, folding and translocation of GCase were assessed in a novel immunofluorescence assay using a GCase antibody that only recognizes the mature lysosomal form of the protein [62]. Overall, these secondary assays provide complementary evidence and allow for triage of hits with the most desirable activity. The combination of the two orthogonal assays also enables deconvolution of allosteric-site-directed GCase enhancers functioning as pure enzymatic activators from those behaving as bona fide PCs driving lysosomal translocation of the protein [54].

### LysoFix-GBA, a fluorescence-quenched substrate, provides direct visualization of GCase function within lysosomes and enables high-content validation of small-molecule GCase enhancers

To implement the LysoFix-GBA assay in high-throughput 384-well microplate format, we first aimed to identify the optimal concentration of the substrate for use in HiBiT-GCase-L444P H4 cells. Automated high-content analysis of subcellular structures provided a readout of integrated LysoFix-GBA spot intensity per cell. LysoFix-GBA demonstrated lysosome-specific activity at concentrations greater than 2.5 μM, which was inhibited by 24 h pre-treatment with the GCase-selective inhibitor AT3375 (**Fig. S6B,C**). At a LysoFix-GBA concentration of 5 μM, HiBiT-GCase-L444P signal was ∼50% of endogenous GCase signal in the *GBA1*-WT H4 line, and minimal background was detected in the *GBA1*-KO H4 line (**Fig. S6D,E**), providing an adequate dynamic range to interrogate modulation of GCase-L444P lysosomal activity. To further characterize the assay, we examined the effects of control compounds NMS-873, bortezomib, isofagomine, NCGC607, and NCGC758 on LysoFix-GBA signal in the HiBiT-GCase-L444P line (**Fig. S6F**). Isofagomine showed an inhibitory LysoFix-GBA response, as expected for an active-site-directed PC; the response to bortezomib was also inhibitory. ERAD modulator NMS-873 showed concordant profiles across the HiBiT and LysoFix-GBA assays, behaving as a consistent enhancer of GCase. Allosteric-site-directed PCs NCGC607 and NCGC758 were both inactive in the LysoFix-GBA assay. Prior studies indicate that NCGC607 and NCGC758 are effective PCs of GCase in different cellular models – macrophages or dopaminergic neurons derived from patients with GD – and given a longer duration of treatment (6 – 21 days) [52, 53]. These findings motivate utilization of the LysoFix-GBA assay as an orthogonal approach to filter hits from the HiBiT assay, allowing for selection of compounds that increase both GCase-L444P levels and lysosomal activity.

Following optimization of the LysoFix-GBA assay in H4 cells, it was deployed on hits arising from the HiBiT-GCase-L444P primary screen, including representatives from each mechanistic cluster (**Fig. 2C**). Pladienolide B, panobinostat, and ARV-825 showed the strongest dose-dependent enhancement of GCase-L444P lysosomal activity (∼3-fold efficacy) after 24 h of compound incubation (**Fig. 3A**). Response to toyocamycin (∼3-fold) peaked at 312.5 nM concentration. Trans-ISRIB recapitulated the nanomolar potency and moderate efficacy (up to 1.8-fold) exhibited in the HiBiT assay after 24 h (**Fig. 3A,B**), but this effect size was increased to more than 5-fold after 72 h of compound incubation (**Fig. S7**). Interestingly, the activatory response to ambroxol peaked at 80-160 nM concentration after both 24 h and 72 h of compound incubation; high concentrations (> 5 μM), which stabilized HiBiT-GCase-L444P protein levels, were found to have an inhibitory effect on enzymatic activity (**Fig. 3A, S7**), as expected for an active-site-directed PC [49]. In contrast, micromolar concentrations of non-inhibitory, allosteric-site-directed PC NCGC326 increased GCase-L444P lysosomal activity up to 2-fold after 24 h (**Fig. 3A,B**) or 72 h (**Fig. S7**) of compound incubation. As an orthogonal dose-response assay, LysoFix-GBA thus validated the pharmacology of several primary screen hits, including PC NCGC326 and some PRs, such as pladienolide B and trans-ISRIB, providing strong support for these compounds increasing levels of functional GCase within lysosomes.

**Figure 3.**
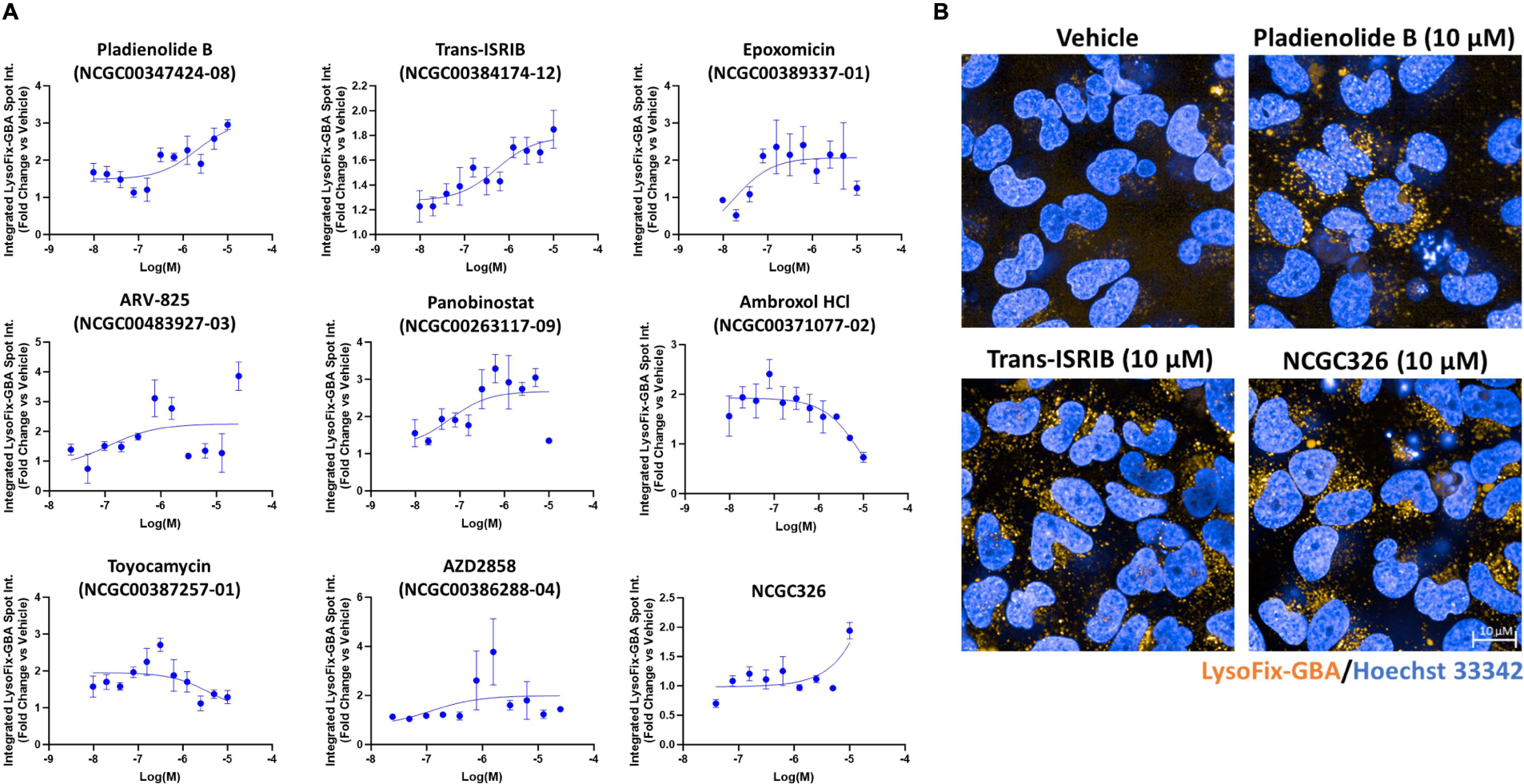
A high-content imaging-based secondary assay using the fluorescence-quenched substrate LysoFix-GBA quantifies GCase activity in the lysosome. Lysosomal activity of GCase was directly visualized using the optimized LysoFix-GBA secondary assay in live HiBiT-GCase-L444P H4 cells treated with vehicle (DMSO) or hit compounds. H4 cells were seeded into 384-well PerkinElmer PhenoPlates (25,000 cells in 40 μL media) and incubated for 24 h. Thereafter, the cells were treated with a titration of compounds for 24 h and then incubated with LysoFix-GBA (5 μM) for 2 h at 37°C and 5% CO_2_. High-content imaging was performed after 15 min of nuclear staining with Hoechst-33342 (1 μg/mL) in Fluorobrite media. (**A**) Select hits from the primary screen were tested in a dose-response titration series. Data are represented as the fold change (compound-treated vs. DMSO-treated) in integrated LysoFix-GBA spot intensity per cell. Dose-response curves were fit using log(agonist) vs. response (three parameters). (Error bars: SEM [*n* = 3 – 6]). (**B**) Representative images are shown for pladienolide B, trans-ISRIB, and NCGC326 at their most effective concentrations in the LysoFix-GBA assay.

### A novel Immunofluorescence-based high-content screening assay allows direct visualization of GCase translocation to the lysosome

We have previously described that the binding of non-inhibitory PCs to GCase can increase its enzymatic activity against specific synthetic substrates [54]. To determine if the increments in GCase activity observed in the LysoFix-GBA assay are due to enzyme activation or an increase in GCase lysosomal translocation, we developed a complementary secondary assay that would provide direct visualization of GCase lysosomal trafficking. We utilized a new monoclonal GCase antibody, hGCase-1/23, which was raised against recombinant GCase (imiglucerase) and is amenable to immunofluorescence studies [62]. Co-staining of GCase with lysosomal marker LAMP1 in 96-well plate format enabled HCS. The assay demonstrated clear lysosomal localization of endogenous GCase in the *GBA1*-WT H4 line and a 77% reduction in signal in the HiBiT-GCase-L444P H4 line (**Fig. 4A,B**). This is consistent with the fact that hGCase-1/23 only recognizes the mature lysosomal form of GCase, not its misfolded, immaturely-glycosylated, ER-retained forms. Through automated high-content analysis, data were quantified as GCase intensity in spots of LAMP1 and normalized to the number of nuclei (**Fig. 4B**). We hypothesized that small molecules which promote proper folding, maturation, and trafficking of GCase-L444P should restore its staining in lysosomes. Indeed, treatment with hit compound pladienolide B (100 nM) caused a 3-fold increase in the co-localization of HiBiT-GCase-L444P with LAMP1 (**Fig. 4A,B**), and the effect was dose-dependent (**Fig. 4C**). In comparison, treatment with NCGC326 (25 μM) or panobinostat (10 μM) for ∼34 h increased GCase/LAMP1 co-localization in the HiBiT-GCase-L444P H4 line by 1.5-fold or 1.6-fold, respectively, relative to DMSO control (**Fig. 4B**). The immunostaining assay thus confirmed that NCGC326 functions as a PC, rather than as a pure GCase activator. Collectively, these results demonstrate the utility of hGCase-1/23 for HCS, enabling visual representation of GCase translocation by PRs and PCs.

**Figure 4.**
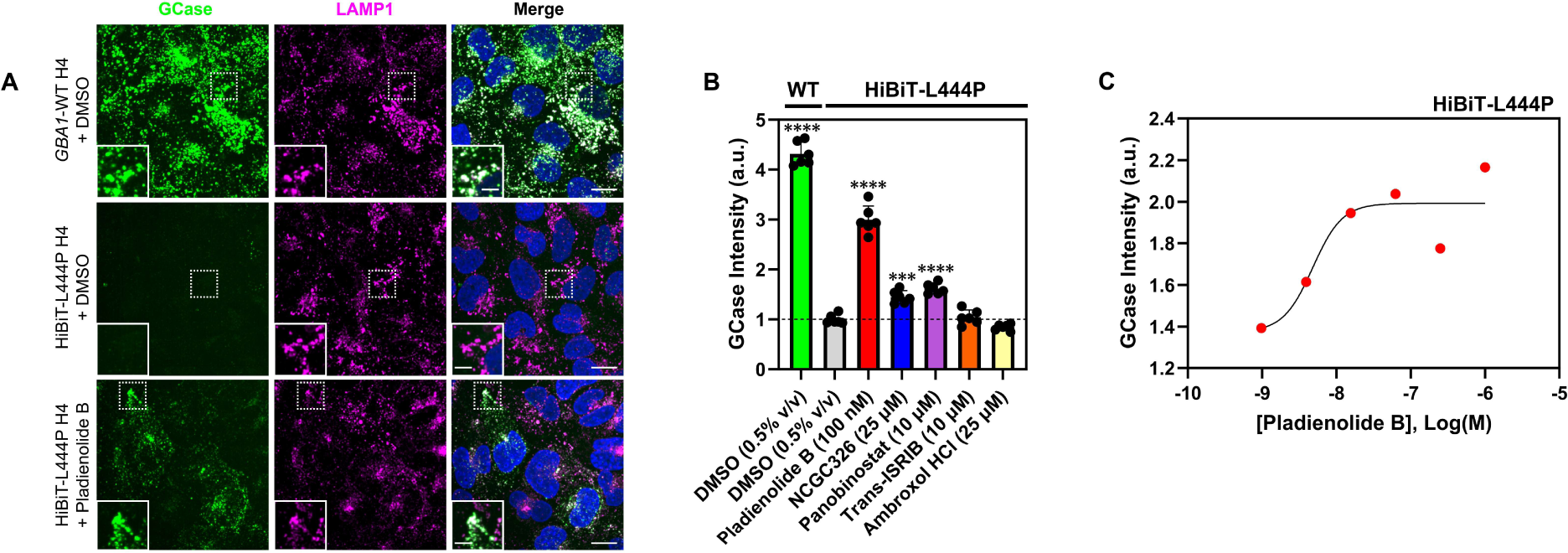
Immunofluorescence secondary assay measures GCase translocation to the lysosome via high-content imaging. HiBiT-GCase-L444P and unedited *GBA1*-WT H4 cells were seeded into 96-well PerkinElmer PhenoPlates and stained for GCase using monoclonal antibody hGCase-1/23 (Alexa Fluor 488), as well as for lysosomal marker LAMP1 (Rhodamine), following treatment with vehicle (DMSO, 0.5% v/v), pladienolide B (100 nM), chaperone NCGC326 (25 μM), panobinostat (10 μM), trans-ISRIB (10 μM), or ambroxol (25 μM) for ∼34 h. Representative images are shown in (**A**), and data are quantified as mean GCase intensity in spots of LAMP1, summed per well, and normalized to cell count, in (**B**). (Error bars: SD [*n* = 6]). (**C**) Dose-response profile of pladienolide B in the GCase/LAMP1 co-localization assay (980 pM – 1 μM; 6-point, 4x dilution) with ∼33 h of treatment. Each data point represents a single replicate that reflects at least 500 cells. Scale bar: 15 μm; Inset scale bar: 5 μm. ***P-value ≤ 0.001 vs. HiBiT-L444P + DMSO; ****P-value ≤ 0.0001 vs. HiBiT-L444P + DMSO.

### Matrix combination screening approach identifies co-formulations of PCs and PRs which synergistically increase HiBiT-GCase-L444P levels

Following the discovery of several PRs that increase GCase-L444P levels through distinct mechanisms of action (**Fig. 2C**), we sought to determine if these PRs would synergize with a PC of GCase [27]. We performed a matrix combinatorial screening assay (**Fig. 5**), in which a titration series of PC NCGC326 was screened in pairwise combination against a titration series of PRs representing different mechanistic classes. Potential synergy was evaluated based on the HiBiT-GCase-L444P response matrix (**Fig. 5A-D**) and Loewe synergy score matrix (**Fig. 5E-H**). The combinations of NCGC326 with PRs ISRIB (**Fig. 5A,E**) or ARV-825 (**Fig. 5D,H**) displayed the greatest synergy across the entire matrix. Pladienolide B and panobinostat also synergized with NCGC326 while inducing great overall fold changes in HiBiT-GCase-L444P levels (**Fig 5B,C**); however, at high concentrations, their synergy (**Fig. 5F,G**) was constrained by strong cytotoxicity (**Fig. S2**).

**Figure 5.**
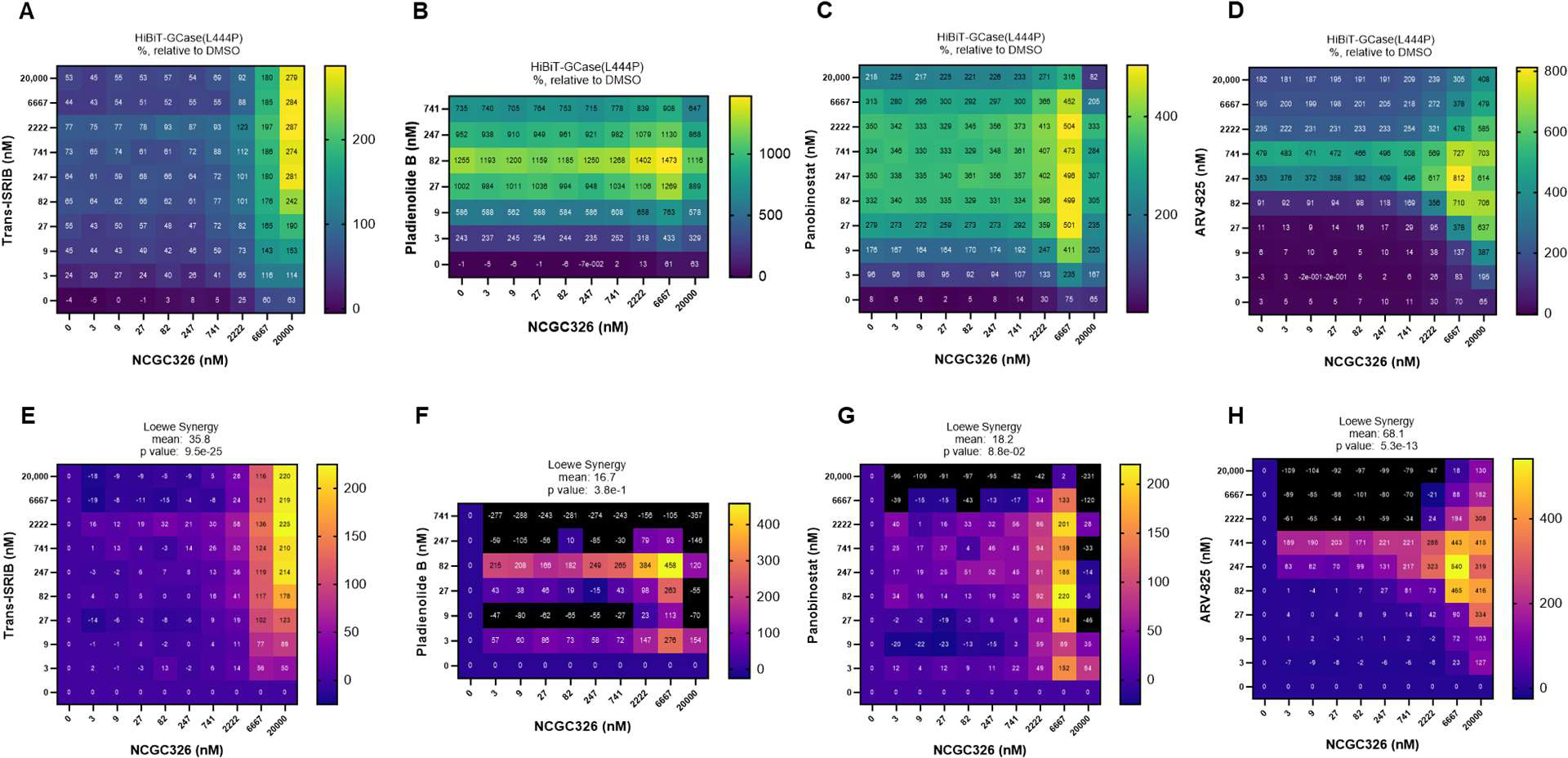
Matrix combination screening approach identifies synergistic co-formulations of a pharmacological chaperone with a proteostasis regulator. HiBiT-GCase-L444P H4 cells were tested in 10 x 10 pairwise dose-response combinatorial matrix format. Cells were treated for 24 h with chaperone NCGC326 in a 9-point titration (3 nM – 20 μM, 3x dilution) against the same 9-point titration of PRs trans-ISRIB (**A, E**), pladienolide B (**B, F**), panobinostat (**C, G**), or ARV-825 (**D, H**); the HiBiT-GCase lytic assay was then performed. Luminescence response values were normalized to intraplate DMSO-treated controls, such that 100% efficacy reflects a doubling of HiBiT-GCase levels. Synergy was evaluated based on the dose-response matrix (**A-D**) and the Loewe synergy score (**E-H**). In general, negative, zero, and positive synergy scores indicate antagonistic, additive, and synergistic interactions, respectively, between drugs. *n* = 3.

### Selected small molecules successfully rescue the metabolic defect of Gaucher disease in HiBiT-GCase-L444P H4 cells

GD is a metabolic disorder that leads to cellular accumulation of the toxic glycosphingolipid species GluCer and GluSph, which are substrates of GCase. To furnish a physiologically-relevant functional endpoint for our drug discovery pipeline, we hypothesized that small molecules which promote GCase-L444P folding, trafficking, and lysosomal activity would correct the biochemical defect contributing to substrate accumulation. We treated HiBiT-GCase-L444P H4 cells for 3 days (starting at 60-70% confluence) with either DMSO (0.3% v/v) or hit compounds ambroxol (156.25 nM), trans-ISRIB (1.25 μM), and NCGC326 (25 μM), as well as the combination of NCGC326 with trans-ISRIB; dose selection was based on 72 h testing with LysoFix-GBA (**Fig. S7**). Levels of GluSph (**Fig. 6**) were then quantitated by supercritical fluid chromatography (SFC) separation coupled with tandem mass spectrometry (MS/MS) detection (SFC-MS/MS) [15, 63]. Impaired clearance of GluSph resulted in 3-fold accumulation of the substrate in HiBiT-GCase-L444P H4 cells, relative to the HiBiT-GCase-WT line, providing a moderate window to interrogate the effect of hit compounds (**Fig. 6A**). NCGC326, as a single agent or in combination with trans-ISRIB, completely reversed lipid accumulation in HiBiT-GCase-L444P H4 cells, to HiBiT-GCase-WT levels (**Fig. 6A**). Ambroxol and trans-ISRIB did not show an effect on lipid accumulation at the concentrations tested (**Fig. 6A**). H4 cells were treated with Pladienolide B (100 nM) for only 48 h (starting at 80-90% confluence), due to the severe cytotoxic effect of the compound (**Fig. S2**); a reduction in GluSph levels was observed (**Fig. 6B**). Collectively, results from the primary screen, both secondary assays, and the lipid functional endpoint support the designation of pladienolide B and NCGC326 as true GCase enhancers in H4 cells.

**Figure 6.**
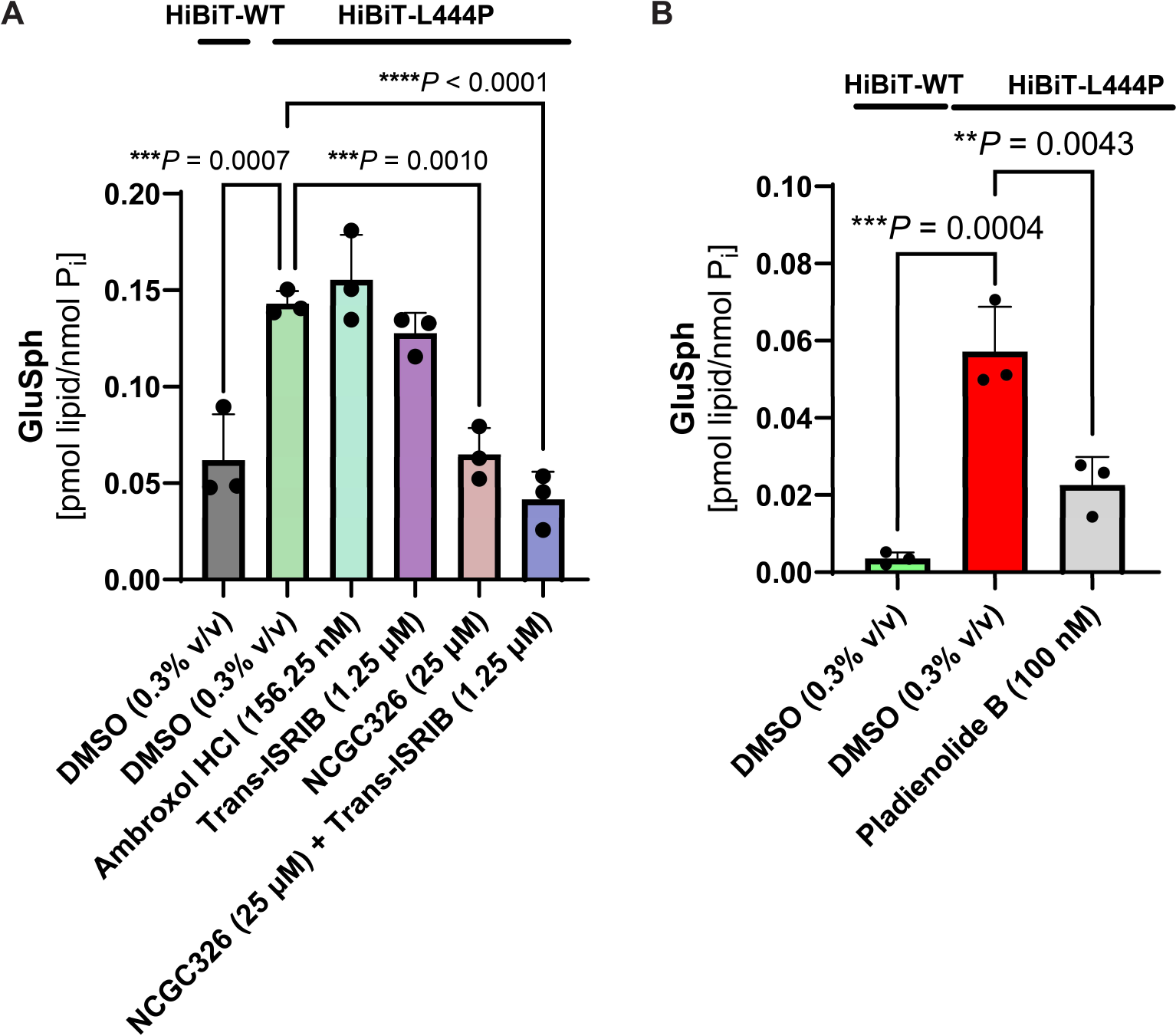
Screening hits reverse glycosphingolipid substrate accumulation in H4 cells. Levels of glucosylsphingosine (GluSph) in frozen cell pellets of HiBiT-GCase-WT and HiBiT-GCase-L444P H4 reporter lines were evaluated by supercritical fluid chromatography (SFC) separation coupled with tandem mass spectrometry (MS/MS) detection and normalized to total cellular inorganic phosphate (P_i_) levels. Cells were treated with vehicle (0.3% v/v DMSO) or hit compounds for 3 days (**A**), except for the cytotoxic hit compound pladienolide B, for which treatment lasted 48 h (**B**). (Error bars: SD [*n* = 3 biological replicates]).

## Discussion

A major challenge in the field of small-molecule GCase enhancers is the lack of adequate cell-based assays amenable to HTS format which can drive the identification, pharmacological validation, and stepwise medicinal chemistry optimization of new chemical matter. To enable rational development of GCase enhancers as a therapeutic strategy, we leveraged an integrated approach, using extensive basic science knowledge surrounding the *GBA1* molecular target to inform the design of physiologically-relevant, translational assays that can accelerate the drug discovery process.

Based on the current paradigm, the pathophysiology of GCase deficiency extends largely from missense mutations in *GBA1* leading to reduced folding efficiency and structural instability of the protein and, consequently, its increased targeting for ERAD. Importantly, these missense-mutant proteins could be catalytically competent in the lysosomal environment, where low pH and high substrate concentration generate a stabilizing milieu [20, 21], but their inability to bypass the ER quality control system creates an obstructive trafficking defect. In particular, the L444P variant of GCase suffers from a perturbed hydrophobic core on a non-catalytic domain [25], which causes overzealous ERAD [19] and correlates with a clinically-severe phenotype [24, 83]. ERAD is the rate-limiting step in the folding of mutant GCase [72], as it controls the reservoir of folding intermediates retained in the ER. The ERAD machinery is part of the larger proteostasis network [84], which also consists of proteinaceous molecular chaperones that promote protein folding, as well as stress-responsive signaling pathways, such as the UPR, heat shock response (HSR), and integrated stress response (ISR), that provide dynamic regulation. Aging-related decline in the robustness and capacity of the proteostasis network escalates protein misfolding and partially underlies the development of neurodegenerative diseases, such as PD [32, 85]. The proteostasis network represents a rational entry point for stabilization of mutant GCase levels, either through enhanced folding mediated by molecular chaperones or through escape from ERAD [21, 72]. Its modification can be achieved with small-molecule PRs, which are a generic biological approach to EET with wide applicability to LSDs and other protein-misfolding diseases [21, 27, 30, 31, 86]. PCs are a separate class of small molecules within EET that are tailored for GCase stabilization through direct binding [42]; their modes of action can include increasing the thermodynamic stability or kinetic accessibility of the native state, or providing dynamic access to a dimer configuration [34–36, 87, 88] that possibly facilitates enzyme trafficking. After PCs and PRs act on mutant GCase to increase its ER export, it passes to the Golgi network, where it undergoes maturation of its glycosylation status before final trafficking to the lysosome. Translocation of GCase from the ER to the lysosome depends upon sorting receptor LIMP-2 [87] and co-chaperone progranulin [89], while its optimal lysosomal activity requires co-factor saposin C [36, 90]. Numerous other genetic modifiers of GCase trafficking and activity likely exist, potentially contributing to the phenotypic heterogeneity observed in GD and the variable penetrance of *GBA1*-PD [91, 92].

Most prior efforts to identify GCase PCs have failed to incorporate these cellular complexities in the discovery phase, relying instead on biochemical assays with purified protein or homogenates [39, 40, 51, 88, 93–95]. Such assays do not align with the true function of PCs, which is salvage of misfolded protein from ERAD, nor do they allow for the identification of PRs, which indirectly regulate GCase through their effects on distinct molecular pathways. Furthermore, cell-free biochemical assays neglect factors such as membrane permeability, intracellular bioavailability, or cytotoxicity. Targeting GCase enhancement in live cells is thus better suited to identify physiologically-relevant small-molecule therapeutics. To this end, we assembled a toolkit of complementary, high-throughput cellular assay modalities. Our novel HiBiT-GCase primary assay evaluates stabilization of GCase protein levels with a luminescence-based readout in 1536-well plate format, enabling qHTS. Two orthogonal, high-content imaging-based secondary assays were then used to triage hits for follow up; these include a 384-well live-cell assay that measures GCase lysosomal activity using the fluorescence-quenched substrate LysoFix-GBA [61] and an immunofluorescence assay that demonstrates direct localization of GCase to the lysosome using the antibody hGCase-1/23 [62]. Finally, reversal of glycosphingolipid substrate accumulation was evaluated by SFC-MS/MS, since this translational endpoint [79] indicates rescue of the biochemical phenotype of GCase deficiency.

The primary assay in this tiered pipeline focuses on stabilization of the severely-misfolded, ERAD-prone L444P variant of GCase in whole cells. To measure GCase protein abundance, we appended a small HiBiT peptide quantitative reporter tag to GCase. HiBiT (1.3 kDa) undergoes high-affinity structural complementation with exogenous LgBiT (18 kDa) in a lytic assay, reconstituting an active luciferase enzyme that affords a sensitive readout. This method of tagging was selected over alternative, larger-size reporters, such as HaloTag (33 kDa) and GFP (27 kDa), as we hypothesized it would be less likely to perturb protein-protein interactions involved in the degradation or trafficking of GCase [96]. Our results indicate that an N-terminal HiBiT tag has negligible influence on GCase maturation, trafficking, and function. To be considered reliable, a quantitative reporter must represent the target biology with high fidelity. Previous studies indicate that stable integration of a HiBiT reporter, via gene editing, more faithfully maintains biological function compared to plasmid-based overexpression systems [97]. Moreover, overexpression of proteins can have mechanistic consequences, including overload of protein quality-control machinery, promiscuous interactions, and stochiometric imbalance [98], which could generate artifacts. Given these considerations, we utilized TALEN-mediated gene editing [66, 67] to stably incorporate a single copy of the HiBiT-tagged *GBA1* variant into an intragenic safe-harbor locus within a *GBA1*-KO background, as affirmed by ddPCR of the selected clones; this workflow constrained *GBA1* expression to a level comparable with the endogenous alleles in the hypertriploid *GBA1*-WT H4 cell line.

When coupled with titration-based qHTS on a fully-integrated robotic screening platform, the HiBiT-GCase-L444P assay generates high-quality pharmacological data, in the form of efficacy and potency values extracted from concentration-response curves [50]. We deployed the assay in a screen of 10,779 small molecules within the NCATS bioactive collections, which include approved drugs, investigational agents, and annotated tool compounds [64]. The results provided prospective validation of the ability of the HiBiT-GCase-L444P assay to detect relevant chemical matter. Ambroxol, a known active-site-directed PC [19, 43, 69, 99] currently being evaluated in a multi-center Phase 3A clinical trial for patients with PD and known *GBA1* status (NCT05778617), was identified as a highly active compound in the unbiased primary screen. Furthermore, the screen identified 32 compounds with efficacy > 300% (4-fold higher than DMSO), of which 27 (84%) were annotated as HDAC inhibitors. As PRs, HDAC inhibitors can reprogram the proteostasis network by causing post-translational hyper-acetylation of histones, transcription factors, and molecular chaperones [73, 100]. This class of small molecules has been investigated in the context of numerous protein-misfolding diseases, including cystic fibrosis [100], Niemann-Pick disease [101], Huntington’s disease [102], type 2 diabetes [103], and PD [104]. The compounds have also been previously reported as enhancers of mutant GCase that might be developed as GD therapeutics [73, 74]. For example, the HDAC inhibitor vorinostat increased the half-life of L444P-mutant GCase by reducing its targeting for ERAD through the molecular chaperone HSP90β [73, 74]. Notably, we have not defined the precise contributions of transcriptional versus post-translational effects of the HDAC inhibitors on GCase in the H4 model system.

The HiBiT primary screen was also able to rank order analogs of in-house lead PCs NCGC607 and NCGC758, indicating its value as a phenotypically-relevant, cell-based tool for lead optimization via medicinal chemistry efforts. Importantly, NCGC326 was identified as a novel analog of NCGC607 with a nearly 3-fold improvement in efficacy and similar potency. While NCGC758 was inactive as a stabilizer of L444P-mutant GCase, its analog, LTI-291 / BIA 28-6156 [78, 79], a compound currently being evaluated in a Phase 2 clinical trial for patients with *GBA1*-PD (NCT05819359), was detected as a hit with similar efficacy to NCGC607. Collectively, these results underscore the reliability of the HiBiT-GCase-L444P assay, its ability to detect both PCs and PRs, and its utility as a workhorse for lead discovery and development.

In addition to validating the high-throughput assay, our primary screen of annotated, chemogenomic libraries enables both drug repositioning and hypothesis generation, which can guide the identification of novel molecular targets regulating the *GBA1* pathway. Analysis of the target profiles of 140 confirmed hit compounds revealed 30 enriched molecular targets, which mainly represented epigenetic regulators, such as HDACs 1-11, NCOR1/2, bromodomain-containing proteins (BRD2, BRD4, BRDT), CHD4, and SIRT6 (**Fig. S3**). The analysis also implicated the Wnt/β-catenin signaling pathway, which is known to be impaired under GCase deficiency [105]. Five confirmed hits inhibited GSK3β, a negative Wnt regulator, while another hit (SKL2001) directly activated Wnt signaling. In agreement, pharmacological Wnt activation was previously shown to rescue defects in dopaminergic neurogenesis [105] and bone matrix deposition [106] in iPSCs derived from patients with GD. Across the identified molecular targets, the top enriched pathway was NOTCH signaling, which has also been previously linked to *GBA1*-PD [107].

Two of the most potent hits identified in our screen were pladienolide B (EC_50_ = 9.4 nM) and trans-ISRIB (EC_50_ = 11.9 nM). Pladienolide B is an inhibitor of the splicing factor SF3B1 [108] that causes production of alternatively-spliced transcripts [109]. While this molecular target has translational potential [109], its mechanistic linkage to GCase stabilization is unclear. Prior studies indicate that low levels of pladienolide B enhance protein-folding capacity by modulating the levels of molecular chaperones [110]. As a PR, ISRIB inhibits the ISR, which responds to the presence of misfolded proteins by reducing global protein synthesis rates [111]. Interestingly, ISRIB inhibits chronic, low-grade ISR activation but spares acute, cytoprotective ISR activation, explaining its lack of overt toxicity *in vivo* [112]. The potency with which ISRIB modifies its target (IC_50_ = 5 nM [76]) aligns with the value determined in our primary screen. Furthermore, its favorable safety profile combined with its cognitive memory-enhancing effects [76] and its demonstrated efficacy in models of neurodegenerative disease, such as prion disease [113] and Alzheimer’s disease [114], make it an attractive therapeutic candidate for *GBA1*-PD. Recently, ISRIB was derivatized into the investigational drug ABBV-CLS-7262, which is under clinical trials for Amyotrophic Lateral Sclerosis (Phase 2/3; NCT05740813) and Vanishing White Matter disease (Phase 1B/2; NCT05757141). Based on their intriguing mechanisms of action, pladienolide B and ISRIB represent high-priority hits for subsequent validation.

Owing to their distinct mechanisms of action, PCs and PRs are candidates for co-formulation into a synergistic combination therapy [27]. This approach is reminiscent of the fixed-dose triple combination therapy used to treat protein misfolding in cystic fibrosis, which involves co-administration of two orthogonal PCs with a CFTR channel gating potentiator [115]. We hypothesized that PRs would increase the population of folded GCase-L444P in the ER, upon which PCs could then act, thus synergistically installing an export-permissive, corrective environment in the ER [27]. To facilitate design of a highly potent and efficacious combination therapy, we deployed a matrix combination screening approach [116, 117] using our HiBiT-GCase-L444P qHTS platform. The combinations of PC NCGC326 with PRs trans-ISRIB or ARV-825 displayed the greatest synergy. Given these results, the widespread deployment of HiBiT-driven matrix combination screening may provide a general strategy to identify combination therapies for other LSDs and loss-of-function protein-misfolding diseases.

Since our drug discovery pipeline addresses GD as a protein-misfolding disease, focusing on defects in GCase folding, trafficking, and lysosomal activity, complementary evidence must be provided at the level of the biochemical phenotype, which is most proximal to disease pathology. Increments in GCase activity as measured by hydrolysis of a synthetic fluorescent substrate, like LysoFix-GBA, might not correlate with improvement in hydrolysis of the natural substrates, GluCer and GluSph [54, 88]. GluCer is the primary storage product in GD and the major glycolipid substrate of GCase; GluSph is the minor glycolipid substrate, being degraded with a ∼100-fold lower turnover number, and its accumulation is thought to be largely driven by deacylation of GluCer by acid ceramidase [118]. Since GluSph is almost undetectable in normal tissues, it represents a sensitive and specific biomarker for GD and its response to therapy [119]. Pathologically, GluSph is neurotoxic [120], possibly as a result of its amphipathic nature and effects on lysosomal membrane permeabilization [121], and it accumulates up to 1,000-fold in the brains of patients with neuronopathic GD [122]. Collectively, these features render GluSph a suitable functional endpoint for evaluation of candidate lead compounds. Given that our HiBiT-GCase-L444P H4 cell model exhibited 3-fold accumulation of GluSph relative to the HiBiT-GCase-WT line, we tested whether the GCase stabilizers discovered from our pipeline could rescue the lipid phenotype. NCGC326 completely reversed the biochemical defect, underscoring the capacity of our pipeline to identify physiologically-relevant chemical matter. Interestingly, ambroxol and trans-ISRIB did not lower GluSph levels, despite enhancing GCase-L444P lysosomal activity at the concentration tested, demanding further investigation.

In this work, we develop and deploy a suite of novel, high-throughput-amenable, quantitative assay technologies that can further the development of small-molecule therapeutic strategies to enhance GCase. Despite the strengths of our assay technologies, the study has some limitations. The chief liability of our HiBiT-based primary assay is that the reporter tag was not appended to the endogenous *GBA1* allele. Rather, exogeneous HiBiT-tagged *GBA1* is constitutively expressed from cDNA integrated into a safe-harbor site within a *GBA1*-KO background; therefore, the model is geared to capture post-translational dynamics and is inadequate to identify GCase stabilizers acting through direct epigenetic or transcriptional regulation from the native locus [123, 124]. Moreover, the HiBiT assay was executed in an immortalized human cell line (H4 neuroglioma), which may feature differential wiring or capacity of the proteostasis network relative to patient cells. To begin to address this limitation, our secondary assays should be adapted to patient-derived neurons for hit validation [81]. Future experiments would benefit from assay technologies enabling measurement of glycosphingolipid substrate levels in high-throughput fashion, including in cell models with greater fold lipid accumulation. Our matrix combination screening approach focused on synergistic combinations of a single PC with select PRs; however, it is possible that dissimilar PRs acting on different modules in the proteostasis network could also exhibit synergy [72]. This would be better explored with an all-versus-all matrix design [117, 125]. Finally, while the small molecules identified in this work may hold therapeutic potential for *GBA1*-PD, or even sporadic PD [3, 14, 15], our pipeline does not feature assays for quantitation of a PD phenotype, such as α-synuclein aggregation [52].

In conclusion, we integrated basic science knowledge surrounding the *GBA1* target with translational expertise to construct a multi-level drug discovery pipeline for GD and *GBA1*-PD. Our approach identified small molecules that increase GCase-L444P protein levels (HiBiT-GCase assay), lysosomal activity (LysoFix-GBA assay), and lysosomal translocation (hGCase-1/23 immunofluorescence assay). These include a PR with a novel mechanism of action, pladienolide B, and an improved derivative of an existing PC, NCGC326. The latter also reversed GluSph accumulation in the H4 model (SFC-MS/MS functional endpoint). Finally, we used a combinatorial matrix screening approach to identify synergistic actions of PRs with a PC in enhancing GCase-L444P levels. In future work, the HiBiT-GCase-L444P assay will be deployed on a diversity library to identify PCs and PRs with new chemical matter. The hits identified in this screen and validated in orthogonal assays should be prioritized for further investigation, with the goal of providing a potent and efficacious therapeutic for patients with neuronopathic GD or *GBA1*-PD.

## Materials and Methods

### Plasmid construction

The *CLYBL*-targeting pC13N-iCAG.copGFP vector (Addgene #66578) [66] was used as a backbone to generate three constructs encoding different N-terminal-HiBiT-*GBA1* variants (WT, N370S, L444P) under the control of a constitutive chimeric CAG promoter. A custom gene block (GENEWIZ) encoding the *GBA1* signaling peptide, HiBiT tag, and Gly/Ser (GS) linker was inserted into the pC13N-iCAG.copGFP vector through BsrGI and MluI digestion and Hi-T4™ ligation (Cat.#M2622; New England Biolabs), thus replacing copGFP and generating the pC13N_N-HiBiT plasmid. *GBA1* transgenes were PCR-amplified from in-house cDNA (using the following 5’ to 3’ primer sequences: Forward: TGGGAATTCTGGTGGAGGATCCgcccgcccctgc; Reverse: acctgaggagtgaattcacgcgttcactggcgacgccac) with Q5® Hot Start High-Fidelity master mix (Cat.#M0494, New England Biolabs) and purified by DNA gel electrophoresis. The pC13N_N-HiBiT vector was prepared to receive the transgene inserts through BamHI and MluI digestion. The linearized vector was assembled with the transgene amplicons via NEBuilder® HiFi DNA Assembly (Cat.#E5520; New England Biolabs), thus generating the three pC13N_N-HiBiT-*GBA1* plasmids. The three ligation products were separately transformed into One Shot™ TOP10 Chemically Competent *E. coli* (Cat.#C404010; Invitrogen), which underwent selection with Kanamycin. A single colony from each plate was then grown in liquid bacterial culture with Kanamycin; the plasmid DNA was amplified using a NucleoBond Xtra Midi kit (Cat.#740410; Macherey-Nagel) and the sequence was verified with Sanger sequencing (GENEWIZ).

### H4 cell culture and transfection

The H4 human neuroglioma cell line (HTB-148) was obtained from American Type Culture Collection (ATCC). An H4 *GBA1* loss-of-function (*GBA1*-KO) clonal cell line was generated through zinc-finger nuclease-mediated targeted integration of gene disruption cassettes into exon 4 of *GBA1* alleles, as previously described [62]. H4 cells were cultured in DMEM + GlutaMAX^TM^ with high glucose (Cat.#10566-016; Gibco) supplemented with 10% heat-inactivated fetal bovine serum and 1 mM sodium pyruvate (Cat.#11360-070; Gibco), with or without 100 U/mL Penicillin-Streptomycin (Cat.#15140-122; Gibco). Cell cultures were maintained in a humidified incubator (37°C, 5% CO_2_) and routinely tested negative for mycoplasma contamination.

HiBiT-*GBA1* constructs were transfected into the H4 *GBA1*-KO cell line using Lipofectamine^TM^ 3000 (Cat.#L3000001; Invitrogen), following the manufacturer’s protocol, with Opti-MEM™ Reduced Serum Medium (Cat.#31985062; Gibco) used for dilution.

### Stable integration of HiBiT-GBA1 transgene into H4 GBA1-knockout cell line

HiBiT-*GBA1* transgenes were stably incorporated into the H4 *GBA1*-KO line using TALEN-enhanced integrative gene transfer [66, 67]. Briefly, *GBA1*-KO H4 cells were seeded at an initial density of 800,000 cells/well in a 6-well plate. After 1 day, the cells were transfected with one of three pC13N_N-HiBiT-*GBA1* donor plasmids (WT, N370S, or L444P; 750 ng/well), along with left and right TALENs (pZT-C13-L1: Addgene #62196; pZT-C13-R1: Addgene #62197; 375 ng/well each) targeting the human Citrate Lyase Beta-Like (*CLYBL)* intragenic safe-harbor locus, which is located in intron 2 (between exons 2 and 3). Lipofectamine^TM^ 3000 (Cat.#L3000001; Invitrogen) was used as the transfection reagent, following the manufacturer’s protocol, with Opti-MEM™ Reduced Serum Medium (Cat.#31985062; Gibco) used for dilution. After 24 h, selection with Geneticin™ (600 ng/μL; G418 sulfate; Cat.#10131027; Gibco) was performed for 1 week to enrich for H4 cells expressing the splicing acceptor – T2A self-cleaving peptide (SA-T2A)-linked Neomycin resistance cassette (NeoR/KanR). With this approach, expression of the resistance gene is driven by the active endogenous *CYLBL* promoter; thus, the antibiotic selection enriches for targeted – as opposed to random – integration events [66, 126]. Following antibiotic selection, single clones with stable integration of the HiBiT-*GBA1* construct were isolated via fluorescence-activated cell sorting into a 96-well plate. Individual surviving clones were then expanded, and selected clones underwent quality control via Sanger sequencing (GENEWIZ) and copy number determination.

### Copy number determination assay

The Droplet Digital polymerase chain reaction (ddPCR) copy number determination assay [125, 127] was performed on a QX200 ddPCR system (Bio-Rad Laboratories) at the Center for Cancer Research (CCR) Genomics Core facility (National Cancer Institute, Bethesda, MD). For sample preparation, a ddPCR reaction mix was prepared which consisted of 1.1 µL of 20X FAM-conjugated, HiBiT-directed target probe (Unique Assay ID: dCNS206145924; MIQE Context: ACAGGATTGCTTCTACTTCAGGCAGTGTCGTGGGCATCAGGTATGGTGAGCGGCTGGCGG CTGTTCAAGAAGATTAGCGGGAGCTCCGGTGGCTCGAGCGGTGGGAATTCTGGTGGAGGA TCC), 1.1 µL of 20X HEX-conjugated reference probe (RPP30; Unique Assay ID: dHsaCP1000485; MIQE Context: ATGAGGAACCTGAAACTTCATGTTAAGTAACTTGTAAGTGGTAGTGCATAGACTTTAAATCAGGCAGACTGACACTAGAGTTCACATTCATAACCACTCCTCAAATGTCCTCCTACTCTTGAC), 1.1 µL of HindIII-HF (Cat.#R3104S; New England Biolabs) diluted with CutSmart® Buffer (Cat.#B7204S; New England Biolabs), 7.7 µL (50 ng) of DNA sample, and 11 µL of 2X ddPCR Supermix for Probes (No dUTP). A plate loaded with 22 µL/well of total sample mixture was placed in an Automated Droplet Generator, after which it was heat sealed with a foil sheet and transferred to a C1000 Touch^TM^ Thermal Cycler. PCR was conducted with a ramp rate of 2°C/s for 1 cycle of enzyme activation (95°C for 10 min), 40 cycles of denaturation (94°C for 30 sec) and annealing/extension (60°C for 1 min), and 1 cycle of enzyme deactivation (98°C for 10 min), after which samples were held at 4°C. The plate was read on a QX200 Droplet Reader, and data were analyzed with QuantaSoft Analysis Pro software (Bio-Rad Laboratories). The results of the ddPCR copy number determination assay informed selection of clones with stable integration of 1 copy of HiBiT-*GBA1*, which was indicated by a FAM/HEX ratio of ∼0.5.

### Western blotting

Cell pellets were lysed in 1% Triton X-100 lysis buffer [1% Triton X-100; 10% glycerol; 150 mM NaCl; 25 mM HEPES pH 7.4; 1 mM EDTA; 1.5 mM MgCl_2_; Pierce^TM^ Protease and Phosphatase Inhibitor Mini Tablets (1 tablet/10 mL; Cat.#A32959)] [62]. The cells were sonicated for 10 quick pulses using a Sonic Dismembrator Model D100 (Fisher Scientific), left standing for 15-30 min on ice, and then centrifuged for 15 min (13,000 x *g*; 4°C). The supernatant containing lysate was collected and stored at −80°C. Protein concentration was evaluated using a Pierce™ BCA Protein Assay Kit (Cat.#23225; Thermo Scientific) by diluting samples in 1% Triton X-100 lysis buffer.

Protein lysates were boiled in loading buffer [1:9 ratio of 2-mercaptoethanol:4x Laemmli Sample Buffer (Cat.#1610747; Bio-Rad Laboratories)] for 10 min. Proteins (40 μg/lane) were resolved on a 4–20% Mini-PROTEAN® TGX Stain-Free™ precast polyacrylamide gel (Cat.#4568094; Bio-Rad Laboratories) and transferred onto a polyvinylidene difluoride (PVDF) membrane via the Trans-Blot Turbo Transfer System (Bio-Rad Laboratories). The membrane was dried, reactivated with methanol, rinsed with water and TBST (1X TBS; 0.1% Tween), and blocked for 1 h at RT [in 1X TBST; 5% (w/v) BSA (Cat.#A-7888; Sigma-Aldrich)]. The membrane was probed overnight (4°C) with primary antibody [0.5 μg/mL in 1X TBST with 2% (w/v) BSA]. Thereafter, it was washed with TBST (3 x 10 min) at RT, incubated with secondary antibody, and washed again with TBST (3 x 10 min, RT). The blot was visualized with SuperSignal West Pico PLUS Chemiluminescent Substrate (Cat.#34580; Thermo Scientific) using the ChemiDoc MP Imaging System (Bio-Rad Laboratories) at 3 sec exposure (with images captured every 6 sec).

The following anti-GBA1 primary antibodies were used: 2E2 (mouse monoclonal; Cat.#H00002629-M01; Abnova; 1:2,000) and R386 (rabbit polyclonal; in-house; 1:10,000). GAPDH was detected with a rabbit polyclonal antibody (Cat.#ab9485; Abcam; 1:15,000). The following goat Horseradish Peroxidase (HRP)-coupled secondary antibodies were used: anti-mouse IgG (Cat.#ab205719; Abcam; 1:10,000) and anti-rabbit IgG (Cat.#ab205718; Abcam; 1:60,000).

### Glycosylation analysis (Endo H and PNGase F assay)

Glycosidase sensitivity analysis was performed as previously described with some modifications [128]. Briefly, protein lysates were prepared in 1% Triton X-100 lysis buffer as described above. For each condition, 40 – 50 µg of protein lysate from each sample was denatured in 1X Glycoprotein Denaturing Buffer (Cat.#B1704S; New England Biolabs) with 20 µL total volume at 100°C for 10 min. The denatured protein lysates were digested with glycosidase enzymes, Endo H_f_ (Cat.#P0703S; New England Biolabs) or PNGase F (Cat.#P0708L; New England Biolabs). For Endo H_f_ digestion, 20 µL of denatured protein lysate was combined with 2.5 µL of 10X GlycoBuffer 3 (Cat.#B1720S; New England Biolabs) and 2.5 µL of Endo H_f_ in a total reaction volume of 25 µL. For PNGase F digestion, 20 µL of denatured protein lysate was combined with 2.6 µL of 10X GlycoBuffer 2 (Cat.#B3704S; New England Biolabs), 2.6 µL of 10% NP-40 (Cat.#B2704S; New England Biolabs), and 1.5 µL of PNGase F in a total reaction volume of 26.7 µL. Undigested control samples were prepared by combining 20 µL of denatured protein lysate with 5 µL of water. The reactions were incubated at 37°C for 1.5 h, after which Western blots were run to examine the digestions. The Endo H-sensitive fraction was designated as immature, ER-retained GCase, while the Endo H-resistant fraction was considered post-ER-localized GCase [24, 128]. Both fractions are responsive to PNGase F treatment.

### AlphaLISA

Amplified Luminescent Proximity Homogeneous Assay (AlphaLISA) was performed using mouse monoclonal antibodies hGCase-1/17 and hGCase-1/23, as previously described [62]. Briefly, 1×10^6^ H4 cells were washed once with PBS and lysed in 80 μL of GCase lysis buffer (0.05 M citric acid, 0.05 M KH_2_PO_4_, 0.05 M K_2_HPO_4_, 0.11 M KCl, 0.01 M NaCl, 0.001 M MgCl_2_, pH 6.0 with 0.1% (v/v) Triton X-100, supplemented with freshly added protease inhibitor).

Samples were diluted 1/5 or 1/10 in 1X Immunoassay buffer (Cat.#AL000F; PerkinElmer). For calibration, a standard curve ranging from 2.4 pM – 1,250 pM of imiglucerase (Genzyme) was generated through 2x serial dilution. 10 μL of samples or standards were incubated for 4 h at RT in the dark with 1 nM of biotinylated hGCase-1/23 and 20 μg/mL of hGCase-1/17-conjugated Acceptor beads (50 :1, beads:antibody). Thereafter, 40 μg/mL of AlphaScreen Streptavidin Donor beads (Cat.#6760002; PerkinElmer) were added and incubated for 1 h at RT in the dark. The assay was performed in a flat white, 384-well OptiPlate (Cat.#6007290, PerkinElmer), which was read on a Tecan Spark plate reader (excitation: 680 nm; emission: 520–620 nm). GCase concentration was calculated based on a sigmoidal non-linear regression of the imiglucerase standard curve and then normalized to total protein concentration.

### GCase activity assay (4-MU-β-Glc)

Enzymatic activity of GCase in protein lysates was determined based on cleavage of the fluorogenic substrate 4-methylumbelliferyl-β-D-glucopyranoside (4-MU-β-Glc) [15]. Briefly, freshly-prepared GCase buffer [McIlvaine buffer (0.2 M Na_2_HPO_4_ and 0.1 M citric acid titrated to pH 5.4); cOmplete™, Mini, EDTA-free Protease Inhibitor Cocktail (Cat.#11836170001; 1 tablet/10 mL); 0.25% (v/v) Triton X-100 (Cat.#T9284; Sigma-Aldrich)] was activated with 0.2% (w/v) sodium taurocholate (Cat.#86339; Sigma-Aldrich). Protein was extracted from H4 cell pellets in GCase buffer through pipetting, at least two 10-sec pulsed bath sonication steps (S-4000, Misonix), and 3 freeze-thaw cycles. Thereafter, the lysates were centrifuged for 15 min (21,000 x *g*; 4°C), and the supernatant containing extracted protein was collected. Protein concentration was measured [Pierce™ BCA Protein Assay Kit (Cat.#23225; Thermo Scientific)], and samples were diluted to 1 mg/mL. Final protein concentration was verified by a second BCA assay and used for data normalization.

The 4-MU-β-D-Glc GCase activity assay was conducted in a Greiner black 384-well plate (Cat.#781209), to which protein samples were added (10 µL/well) and further diluted with GCase buffer (5 µL/well). The plate was sealed, centrifuged, and incubated (37°C) in a shaking plate incubator at 600 rpm for 15 min. After brief centrifugation, 15 µL of assay buffer (2.5 mM 4-MU-β-D-Glc; 0.25% v/v DMF in GCase buffer) was added to each assay well. The plate was sealed, centrifuged, and incubated (37°C) for 60 min, shaking at 450 rpm. After incubation, the plate was centrifuged, and 30 µL of 1 M glycine stop solution (pH 10.5) was added to each well. Fluorescence was top read on a FlexStation® 3 Multi-Mode Microplate Reader (Molecular Devices; 365 nm excitation wavelength; 449 nm emission wavelength; 435 nm cutoff; 6 reads/well). Relative GCase activity was calculated by adjusting for protein concentration, correcting for *GBA1*-KO H4 cell background, and normalizing to *GBA1*-WT signal.

### Immunocytochemistry

For preliminary GCase/LAMP1 co-localization analysis (**Fig. 1C**), H4 cells were seeded into 8-well imaging chambers at a density of 30,000 cells/chamber, 48 h prior to immunostaining. For high-content screening with small molecules, H4 cells were seeded into 96-well, black, optically clear, flat-bottom, tissue-culture treated PhenoPlates (PerkinElmer) at 5,000 cells/well in 200 µL media and treated at ∼50% confluence.

For immunostaining [62], cells were washed once with PBS (pH 7.4) and fixed with 4% paraformaldehyde (20 or 30 min, RT). Fixed cells were washed with PBS (3 x 5 min) and incubated overnight (4°C) with primary antibody in ICC antibody diluent [PBS containing 0.05% or 0.1% saponin (filtered) and 1% BSA]. After primary antibody incubation, cells were washed with PBS (3 x 5 min, at least) and incubated for 1 h (RT) with secondary antibody in antibody diluent. After secondary antibody incubation, cells were washed with PBS (3 x 5 min, at least) and underwent nuclear staining. For Hoechst staining, Hoechst 33342 solution (Cat.#62249; ThermoFisher Scientific) was diluted in PBS to 5 µg/mL and added during the third wash. For DAPI staining, cells were stored at 4°C in PBS with NucBlue™ Fixed Cell ReadyProbes™ Reagent (DAPI) (Cat.#R37606; ThermoFisher Scientific) until imaging. Imaging plates were sealed with a heat-resistant film (Cat.#89496-565; VWR International).

The following primary antibodies were used: GCase (5 µg/mL; hGCase-1/23; mouse monoclonal; Roche [62]); LAMP1 (1:500; D2D11, rabbit monoclonal; Cat.#9091; Cell Signaling Technology). The following secondary antibodies were used: Rhodamine Red™-X (RRX) AffiniPure Donkey Anti-Rabbit IgG (H+L) (1:200; Cat.#711-295-152; Jackson ImmunoResearch Laboratories, Inc.); Alexa Fluor® 488 AffiniPure Donkey Anti-Mouse IgG (H+L) (1:400; Cat.#715-545-150; Jackson ImmunoResearch Laboratories, Inc.).

Regular confocal images were acquired with a Zeiss LSM 880 confocal microscope using a 63x oil objective. High-content imaging was conducted on a PerkinElmer Opera Phenix Plus confocal high-content screening system; images were captured in 27 fields per well with a 63x water, NA 1.15 objective. Image analysis was performed using PerkinElmer Columbus software and algorithms to quantify the sum per well of mean GCase intensity (Alexa 488 channel) in spots of LAMP1 (TRITC channel), which was normalized to number of nuclei (DAPI channel).

### Lipidomic analysis

For initial analysis of glucosylsphingosine (GluSph) levels in frozen pellets of untreated H4 cells, cells were lysed by addition of distilled water followed by protein precipitation with methanol, as described [8, 62]. The dry lipid samples were reconstituted in a mixture of acetonitrile/water 90/10 (v/v) containing 1% DMSO. Analysis was performed via positive ion electrospray LC-MS/MS in multiple reaction-monitoring (MRM) mode, using deuterated compounds as internal standards. A Waters Xevo-TQ-S mass spectrometer connected to a complete Waters Acquity I-class UPLC system was used as previously described, with a mixture of acetonitrile/methanol/water 40/40/20 (v/v/v) as the wash solvent [8, 62]. Samples were analyzed on a BEH glycan amide column (100 × 2.1 mm, 1.7 μm particle size; Waters Corporation, Switzerland) with a flow rate of 0.25 mL/min and an oven temperature of 30°C. Eluent A consisted of 100 mM ammonium acetate, while eluent B was acetonitrile. Glycospecific separation was achieved by isocratic elution with 90% eluent B, followed by a washing step with 10% eluent B and column reconditioning. Each sample was run as five technical replicates with 1×10^6^ cells each. Lipid levels were calculated by linear regression, using the peak area ratio of analyte to internal standard as the response, and expressed as pmol/million cells.

As an endpoint in the drug discovery pipeline, lipids were extracted from frozen cell pellets of compound-or DMSO-treated H4 cells, and levels of GluSph were determined by supercritical fluid chromatography (SFC) separation coupled with tandem mass spectrometry (MS/MS) detection [15, 63, 129]. Cell pellets were washed with 0.15 M NaCl solution (Cat.#46-032-CV; Corning) and stored at −80°C prior to analysis. Extracted lipid samples and synthetic standards were processed on a setup comprising a Waters Acquity SFC/UPC2 Chromatography System and TSQ Quantum Access Max Triple Quadrupole Mass Spectrometers (Thermo Scientific) operating in positive MRM mode and employing a gradient elution. Quantitation required the generation of analyte-specific eight-point calibration curves (analyte/internal standard peak area ratio versus analyte concentration), as described previously [130]. Lipid measurements [pmol/total sample] were normalized to levels of inorganic phosphate [nmol] in each sample, as determined by Bligh & Dyer (B&D) re-extraction [131] of an aliquot from the original total sample extract. Lipid extraction and SFC-MS/MS analysis were performed by the Analytical Unit of the Lipidomics Shared Resource at the Medical University of South Carolina (MUSC) Hollings Cancer Center.

### Microscale thermophoresis

Microscale thermophoresis (MST) experiments were performed using the Monolith NT.Automated instrument (NanoTemper Technologies). Briefly, enzymatically-active recombinant human GCase with a C-terminal 6-His tag (Cat.#7410-GHB-020; Bio-Techne) was purchased as a 0.2 μm filtered solution in 50 mM sodium citrate, pH 5.5 at a concentration of 0.319 mg/mL. The recombinant GCase was further diluted with assay buffer (50 mM sodium citrate, pH 5.5) to 0.160 mg/mL and incubated (RT) for 30 min with 50 nM RED-tris-NTA 2nd Generation dye from the Monolith His-Tag Labeling Kit (Cat.#MO-L018; NanoTemper Technologies). The labeling reaction mixture was centrifuged for 10 min at 4°C and 15,000 x *g* to remove protein aggregates, after which the supernatant was diluted 2.5x with assay buffer. For each compound measured, a 12-point, 2x titration series of compound in DMSO was prepared; 1 μL of compound in DMSO from each point in the titration series was combined with 9 μL of assay buffer and 10 μL of labeled GCase in a 384-well plate, which was left standing for 30 min (RT). The resultant final concentration of labeled GCase was ∼570 nM (in 5% DMSO). MST measurements were conducted with medium MST power, 15% excitation power, red (Pico) excitation color, and 1.5-sec on time using Monolith NT.Automated capillary chips (Cat.#MO-AK002). Data analysis was performed using MO.Affinity Analysis v2.3 software (NanoTemper Technologies), with data fit to a K_d_ model.

### Miniaturization and high-throughput screening of HiBiT-GCase and CellTiter-Glo assays

H4 cells expressing HiBiT-GCase (WT, N370S, or L444P) were grown to 70% confluency and seeded into 1536-well solid white TC-treated microplates (Cat.#782073, Greiner) at a density of 2,000 cells/well in 5 μL media, covered with stainless steel cell culture lids, and incubated for 24 h at 37°C and 5% CO_2_. Cells were then treated with 50 nL of compounds or vehicle (DMSO) via acoustic dispensing (Echo, Beckman Coulter) and incubated for 24 h at 37°C and 5% CO_2_. Assessment of total cellular HiBiT-GCase protein abundance was performed using the Nano-Glo® HiBiT Lytic Detection System (Cat.#N3040, Promega), according to the manufacturer’s protocol. Briefly, the LgBiT Protein (1:100) and the Nano-Glo® HiBiT Lytic Substrate (1:50) were diluted into RT Nano-Glo® HiBiT Lytic Buffer and added at a 1:1 ratio with cell culture media. Cells were incubated with the HiBiT lytic mixture for 10 minutes at RT, and luminescence was measured on a PHERAstar FSX microplate reader with a LUM plus optical module (3600 gain). Settling time and measurement time were selected for optimal precision measurement over speed. Data analysis was performed by comparing the fold change in HiBiT luminescence in compound-treated cells versus vehicle-treated cells. Assessment of cell viability was performed using Promega CellTiter-Glo® Luminescent Cell Viability Assay according to the manufacturer’s recommendation. Briefly, CellTiter-Glo was added at a 1:1 ratio in replicate assay plates with cell culture media. Cells were incubated with the CellTiter-Glo buffer for 10 minutes at RT, and luminescence was measured on a PHERAstar FSX microplate reader using the same settings as the HiBiT assay.

### Primary screen hit selection

Primary screen hits were determined based on a system of curve classes [50], which provides a heuristic measure of data confidence. The amended qHTS curve classification system (CC-v2) was utilized [132]. Curve classes of 1 are complete curves containing two asymptotes and an inflection point, with data confidence decreasing from curve class 1.1 to 1.4. Curve classes of 2 are incomplete curves containing one asymptote and an inflection point, with data confidence decreasing from curve class 2.1 to 2.4. Compounds with a curve class of 5 are inconclusive and were included, so as to not lose any potential hits that could not be classified. For the primary screens, all compounds from CC-v2 1.1-1.4, 2.1-2.4, and 5 were included; further hit selection was applied based on an efficacy cutoff of 40%, and compounds that showed efficacy at a single concentration while also displaying inhibitory effects at a majority of other concentrations were removed (formally defined as an AUC default cutoff of −50%). For the follow-up screens, data were included for all compounds irrespective of curve class, and hit selection was applied using the above cutoffs.

### Target profile and pathway analysis of hit compounds

To gain insight into possible mechanisms of actions for the 140 confirmed follow-up hits, we first built a protein-target profile for each hit compound by collecting target annotations from the NCATS in-house database as well as by querying human target proteins for each compound from five public repositories: ChEMBL [133], DrugBank [134], IUPHAR [135], PharmGKB [136], and PubChem [137]. A bioactivity cutoff of 10 µM or less was used when querying targets from PubChem, ChEMBL, and IUPHAR. Of the 140 hits, target annotations were obtained for 83 compounds while the remaining 57 could not be resolved. Each hit was annotated with at least one target protein (represented by their UniProt accession number), and collectively these 83 hits mapped to 274 unique target proteins, as many compounds target multiple proteins (hit-compounds-targets profile). A protein-target profile was also built for the entire collection of primary screening compounds (background-compounds-targets profile) which collectively mapped to 2773 unique target proteins. Fisher’s exact test was used to conduct a statistical overrepresentation analysis to identify targets that were over-represented in the hit-compounds-targets profile compared to background. The resulting *p*-value from the Fisher’s exact test was corrected via the false discovery rate (FDR) using the Benjamini–Hochberg method. Applying an FDR cutoff of 5% resulted in 30 enriched targets. The known and predicted interaction network of these targets was determined using the STRING database [138]. The UniProt accession number of the enriched targets was then used as input to the Reactome pathway database [139] to identify pathways that are overrepresented by these targets, revealing potential pathways affected by hit compounds.

### Miniaturization and high-throughput screening of LysoFix-GBA secondary assay

H4 cells expressing HiBiT-GCase-L444P were grown to 70% confluency and seeded into 384-well black, optically clear flat-bottom, tissue-culture treated PhenoPlates (PerkinElmer) at a concentration of 25,000 cells/well in 40 μL Fluorobrite media for 24 h at 37°C and 5% CO_2_. Cells were then treated with 50 nL of compounds or vehicle (DMSO) via acoustic dispensing and incubated for either 24 h or 72 h at 37°C and 5% CO_2_. To assess lysosomal GCase activity, LysoFix-GBA was synthesized as previously described [61] and resuspended in DMSO to a stock concentration of 10 mM. Working stocks were prepared by diluting LysoFix-GBA in DMSO. Cells were then treated to desired concentration with 50 nL of the LysoFix-GBA working stock via acoustic dispensing and either imaged immediately for live-cell imaging at 37°C and 5% CO_2_ or incubated for 2 h at 37°C and 5% CO_2_ and imaged after adding Hoechst-33342 prepared in Fluorobrite media (1 μg/mL) for 15 min. Imaging was performed on a PerkinElmer Opera Phenix Plus confocal high-content screening system. Image analysis was performed using PerkinElmer Columbus software and algorithms to quantify the number, intensity, and area of LysoFix-GBA spots per cell indicated by nuclei number. The data were then represented as fold change of integrated LysoFix-GBA spot intensity per cell in compound-treated cells versus vehicle-treated cells. Presented images are contrasted to the highest intensity condition within each experiment.

### Drug synergy evaluation

Synergy of drug combinations was analyzed by the SynergyFinder package [140] using the Loewe additivity model [141] to calculate synergy scores. Each drug concentration combination had three replicates which were used to fit dose-response curves. For compounds NCGC00347424 (pladienolide B), NCGC00389337 (epoxomicin), and NCGC00386288 (AZD2858), the top-three concentrations were removed from analysis, as a dose-response curve could not be fit due to toxicity at higher concentrations. A response matrix and corresponding synergy score matrix were generated for each combination. In general, negative, zero, and positive scores in the synergy matrix indicate antagonistic, additive, and synergistic interactions between drugs, respectively.

### Preparation of N-(4-iodophenyl)-2-(2-((4-iodophenyl)amino)-2-oxoethoxy)benzamide (NCGC00241326)

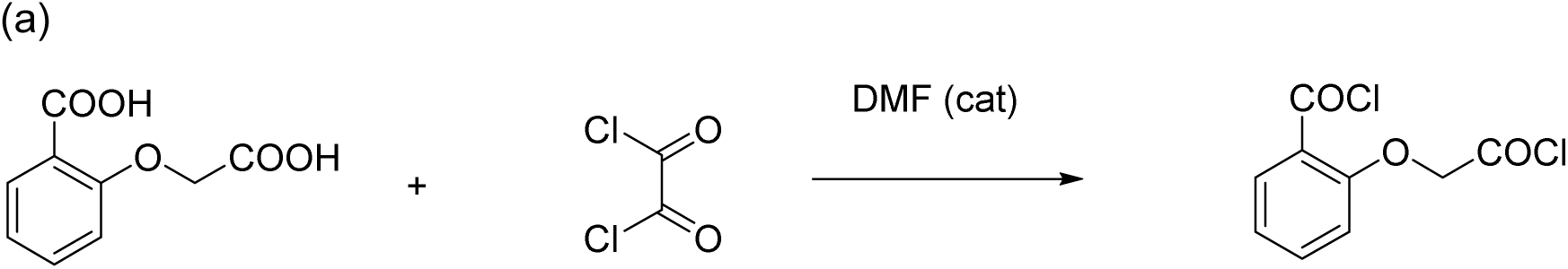

2-(carboxymethoxy)benzoic acid (0.196 g, 1 mmol) and oxalyl dichloride (0.429 mL, 5.00 mmol) in DCM (1 mL) was allowed to stir at RT. One drop of DMF (3.87 µL, 0.050 mmol) was added. The solution was stirred at RT for 30 min. The solution was concentrated. The residue was used as is for the next step.

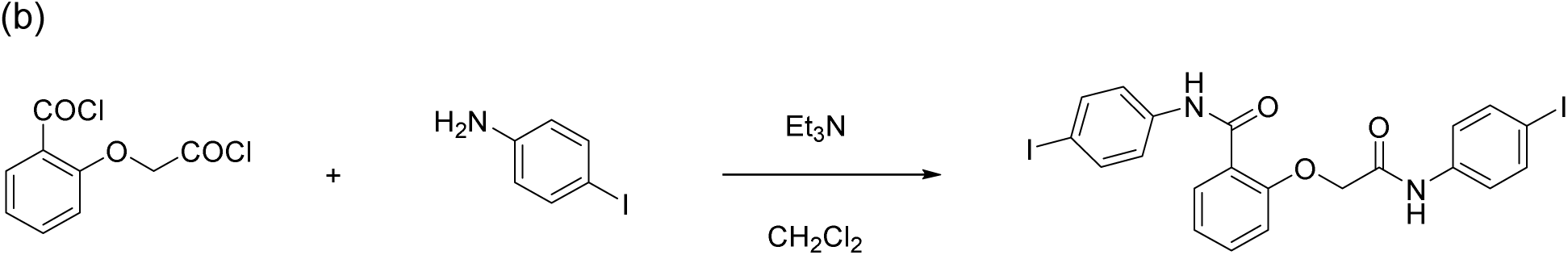

4-iodoaniline (0.482 g, 2.200 mmol) was dissolved in methylene chloride (1 mL). To the solution was added triethylamine (0.348 ml, 2.500 mmol). The solution was stirred at RT. A solution of 2-(2-chloro-2-oxoethoxy)benzoyl chloride (0.233 g, 1 mmol) in methylene chloride (1.000 mL) was added dropwise over a period of 1 min. The solution was stirred at RT for 2 hr. The solution was filtered. The solid was washed with methylene chloride. The solid was dried *in vacuo* to give the desired product, N-(4-iodophenyl)-2-(2-((4-iodophenyl)amino)-2-oxoethoxy)benzamide (117.4 mg, 20% yield). ^1^H NMR (400 MHz, DMSO) δ 10.77 (s, 1H), 10.46 (s, 1H), 7.85 (dd, J = 7.7, 1.8 Hz, 1H), 7.77 – 7.69 (m, 6H), 7.60 (ddd, J = 8.9, 7.4, 1.8 Hz, 1H), 7.54 – 7.48 (m, 2H), 7.28 – 7.23 (m, 1H), 7.20 (td, J = 7.5, 0.9 Hz, 1H), 5.01 (s, 2H). Contains 3.5% methylene chloride. (LC-MS, ESI pos.) Calculated for C_21_H_16_I_2_N_2_O_3_: 599.18 (M + H), Measured: 598.9.

Analytical analysis was performed on an Agilent LC/MS (Agilent Technologies, Santa Clara, CA). A 7-min gradient of 4% to 100% acetonitrile (containing 0.025% trifluoroacetic acid) in water (containing 0.05% trifluoroacetic acid) was used with an 8-min run time at a flow rate of 1.0 mL/min. A Phenomenex Luna C18 column (3 micron, 3 x 75 mm) was used at a temperature of 50°C. Purity determination was performed using an Agilent diode array detector.

### Statistical analysis

Differences between groups were analyzed via either ordinary one-way ANOVA with Tukey’s multiple comparisons test or Brown-Forsythe and Welch ANOVA tests with Dunnett’s T3 multiple comparisons test. Statistical analysis was performed using GraphPad Prism software (version 10.1.2).

## Abbreviations

AlphaLISA: Amplified Luminescent Proximity Homogeneous Assay
BBB: Blood-brain barrier
BET: Bromodomain and extraterminal domain
BRD: Bromodomain-containing protein
BRDT: Bromodomain testis-specific protein
CC: Curve class
CFTR: Cystic fibrosis transmembrane conductance regulator
CHD4: Chromodomain-helicase-DNA-binding protein 4
*CLYBL*: Citrate Lyase Beta-Like
ddPCR: Droplet Digital polymerase chain reaction
DLB: Dementia with Lewy bodies
DMSO: Dimethyl sulfoxide
EET: Enzyme enhancement therapy
ERAD: Endoplasmic reticulum-associated degradation
ERT: Enzyme replacement therapy
GCase: Glucocerebrosidase
GD: Gaucher disease
GluCer: Glucosylceramide
GluSph: Glucosylsphingosine
GSK-3: Glycogen synthase kinase 3
HCS: High-content screening
HDAC: Histone deacetylase
HEAL: Helping to End Addiction Long-term
HSR: Heat shock response
HTS: High-throughput screening
ISR: Integrated stress response
LC: Liquid chromatography
LSD: Lysosomal storage disorder
MS: Mass spectrometry
4-MU-β-Glc: 4-methylumbelliferyl-β-D-glucopyranoside
NCATS: National Center for Advancing Translational Sciences
NCOR: Nuclear receptor co-repressor
NPACT: NCATS Pharmacologically Active Chemical Toolbox
NPC: NCATS Pharmaceutical Collection
PC: Pharmacological chaperone
PD: Parkinson’s disease
PR: Proteostasis regulator
qHTS: Quantitative high-throughput screening
RLU: Relative light units
RT: Room temperature
SAHA: Suberoylanilide hydroxamic acid
SIRT6: Sirtuin 6
SFC: Supercritical fluid chromatography
SMILES: Simplified molecular-input line-entry system
STRING: Search Tool for Retrieval of Interacting Genes/Proteins
TALEN: Transcription activator-like effector nuclease
UPR: Unfolded protein response
VCP: Valosin-containing protein

## Data availability

Final primary screen hits arising from this work are listed in **Table S1**.

## Acknowledgements

This work was supported by the Michael J. Fox Foundation for Parkinson’s Research (MJFF-022588) and the Intramural Research Programs of the National Human Genome Research Institute, National Center for Advancing Translational Sciences, and National Institutes of Health. We thank the NCATS Compound Management and Automation teams for their contributions to this study.

## Supplemental Figures

**Figure S1.**
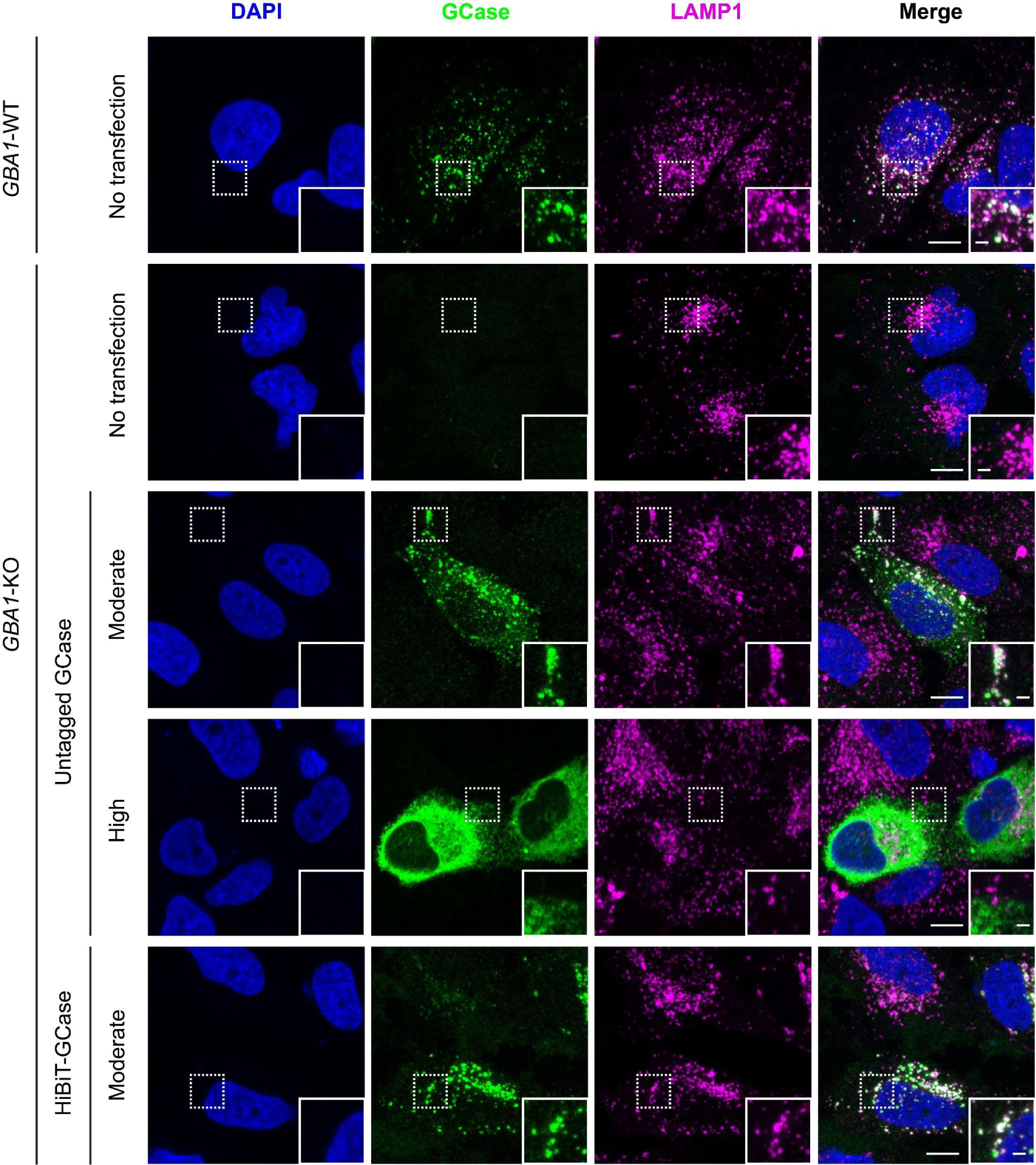
Overexpression of *GBA1* drives mis-localization. *GBA1*-WT or *GBA1*-KO H4 cells were transfected with constructs containing the HiBiT-*GBA1* transgene or untagged *GBA1*. Transfection produced a heterogenous population of cells with either low, moderate, or high levels of GCase expression; the latter perturbed GCase trafficking to the lysosome. GCase was stained with hGCase-1/23 antibody (green), while lysosomes were stained for LAMP1 (magenta).

**Figure S2.**
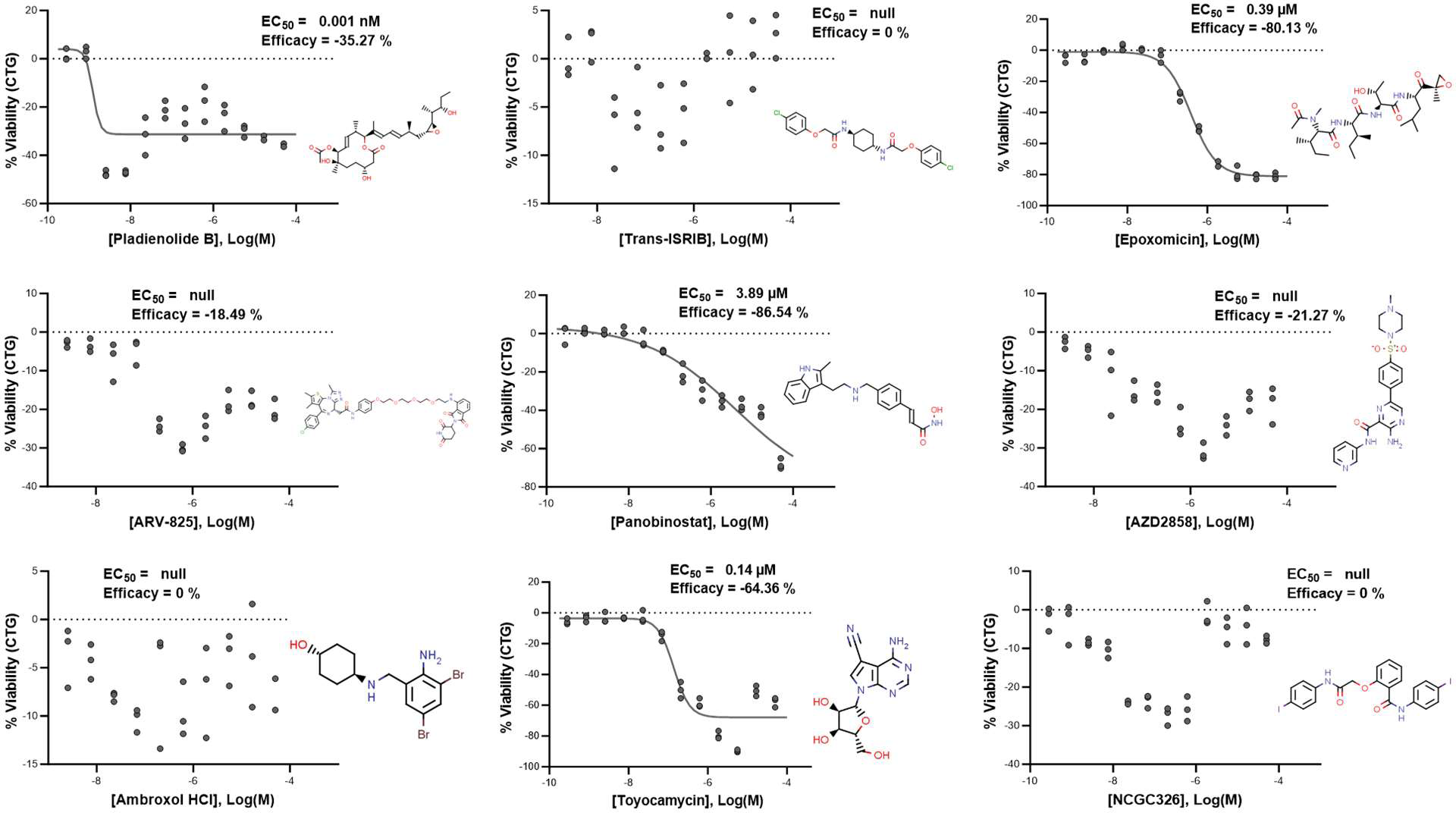
Follow-up cytotoxicity testing of selected primary qHTS hits using CellTiter-Glo assay. HiBiT-GCase-L444P H4 cells were seeded into 1536-well solid white plates (2,000 cells in 5 μL media) for 24 h and incubated with a titration of representative hits from the HiBiT primary screen for 24 h at concentrations ranging from 0.3 nM – 50 μM (12-point, 3x dilution series; *n* = 3). The CellTiter-Glo assay was then performed as described in the methods. Response values (% viability) are based on change in luminescence (RLU) in compound-treated versus DMSO-treated cells.

**Figure S3.**
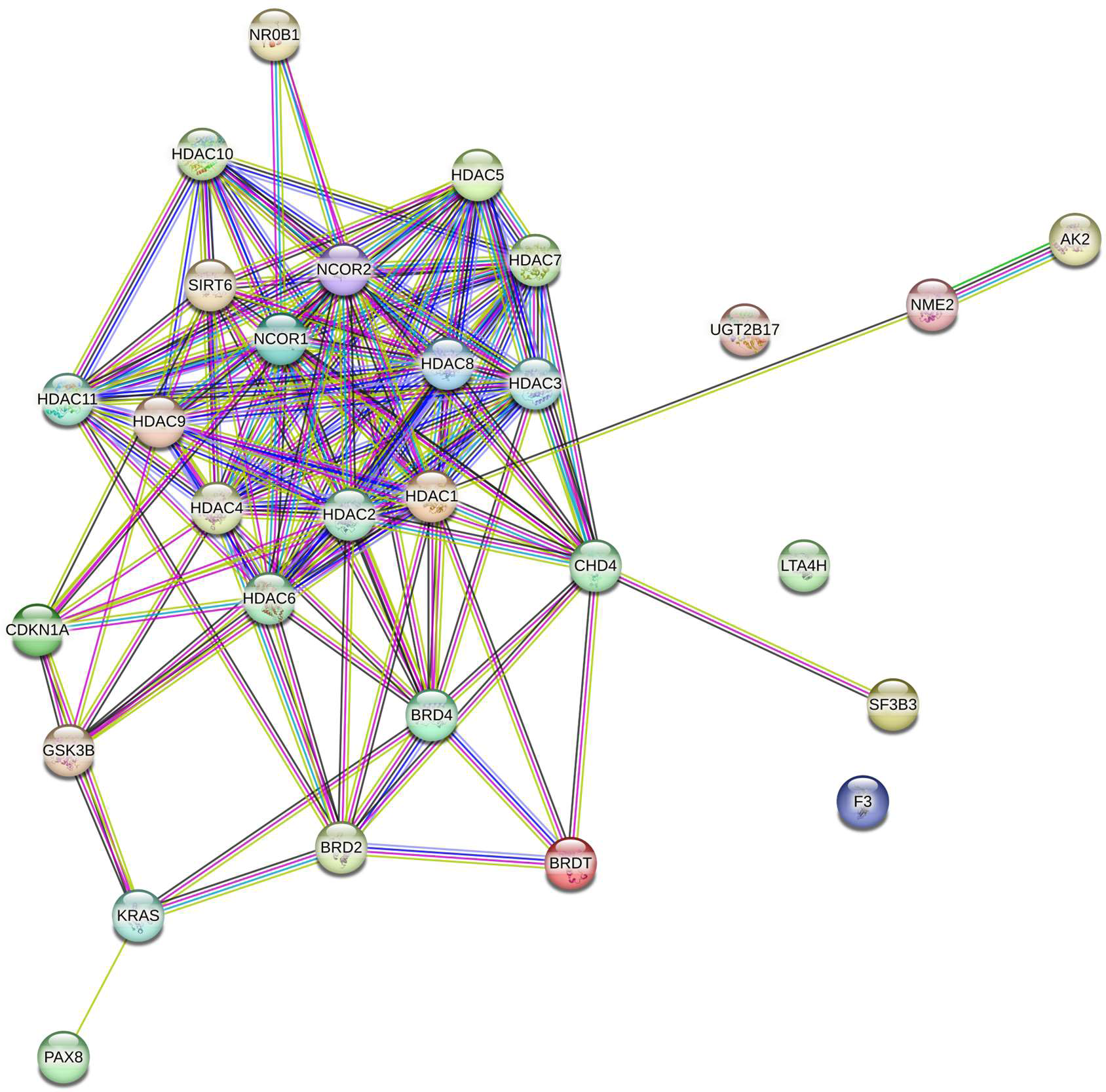
Interaction network of molecular targets of primary qHTS hits. Known and predicted protein-protein interaction network from STRING database for the 30 enriched targets identified from target profiles of final hit compounds. **Table S2** lists the pathways in the Reactome database that are overrepresented by these enriched targets, revealing potential pathways affected by the hit compounds.

**Figure S4.**
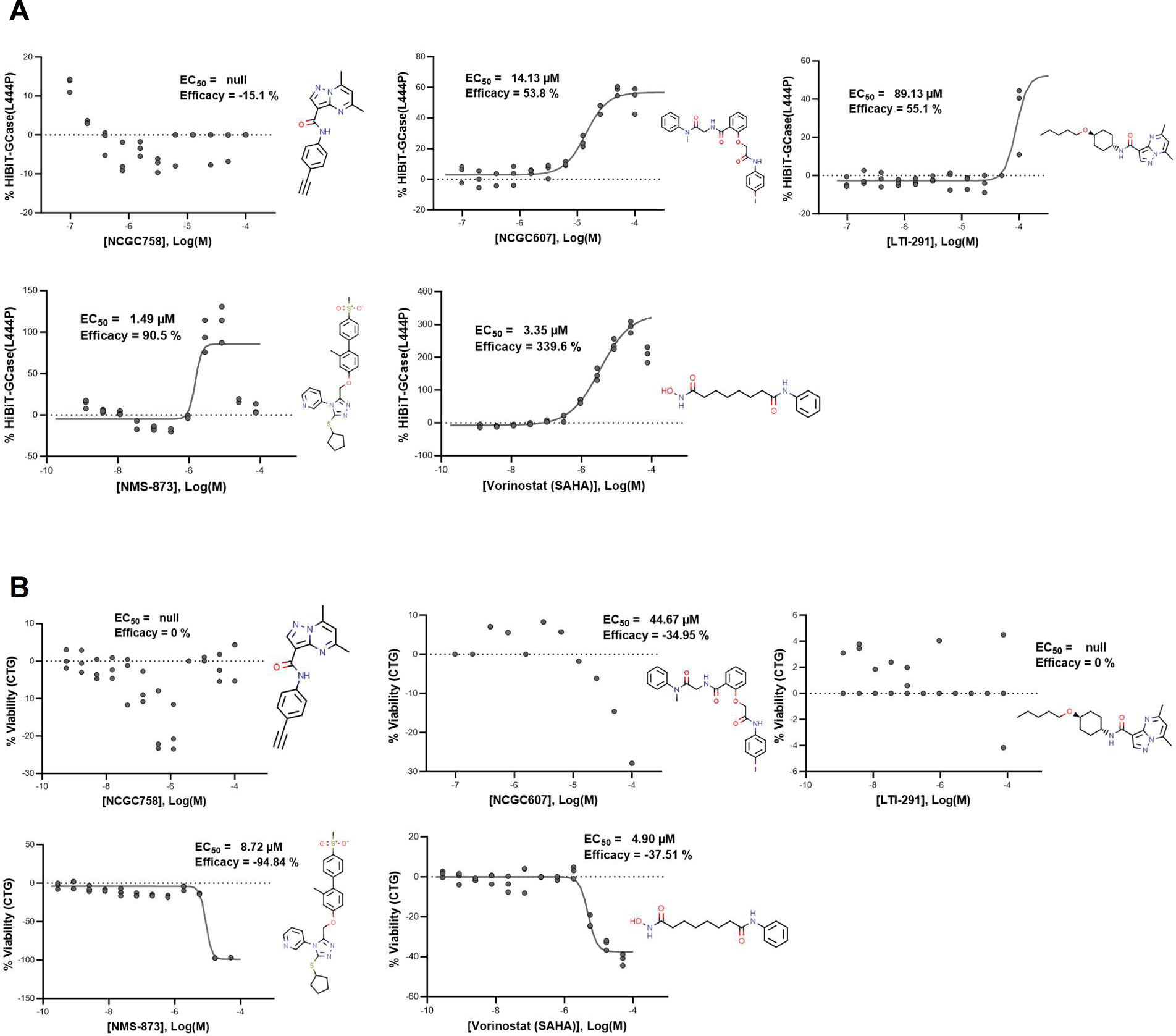
Primary qHTS curves and cytotoxicity testing for additional compounds. HiBiT-GCase-L444P H4 cells were seeded into 1536-well solid white plates (2,000 cells in 5 μL media) for 24 h and treated with a titration of NCGC758, NCGC607, LTI-291, NMS-873, and vorinostat (SAHA) for 24 h, after which either the HiBiT assay (**A**) or CellTiter-Glo assay (**B**) was performed as described in the methods. Data are from three independent replicates.

**Figure S5.**
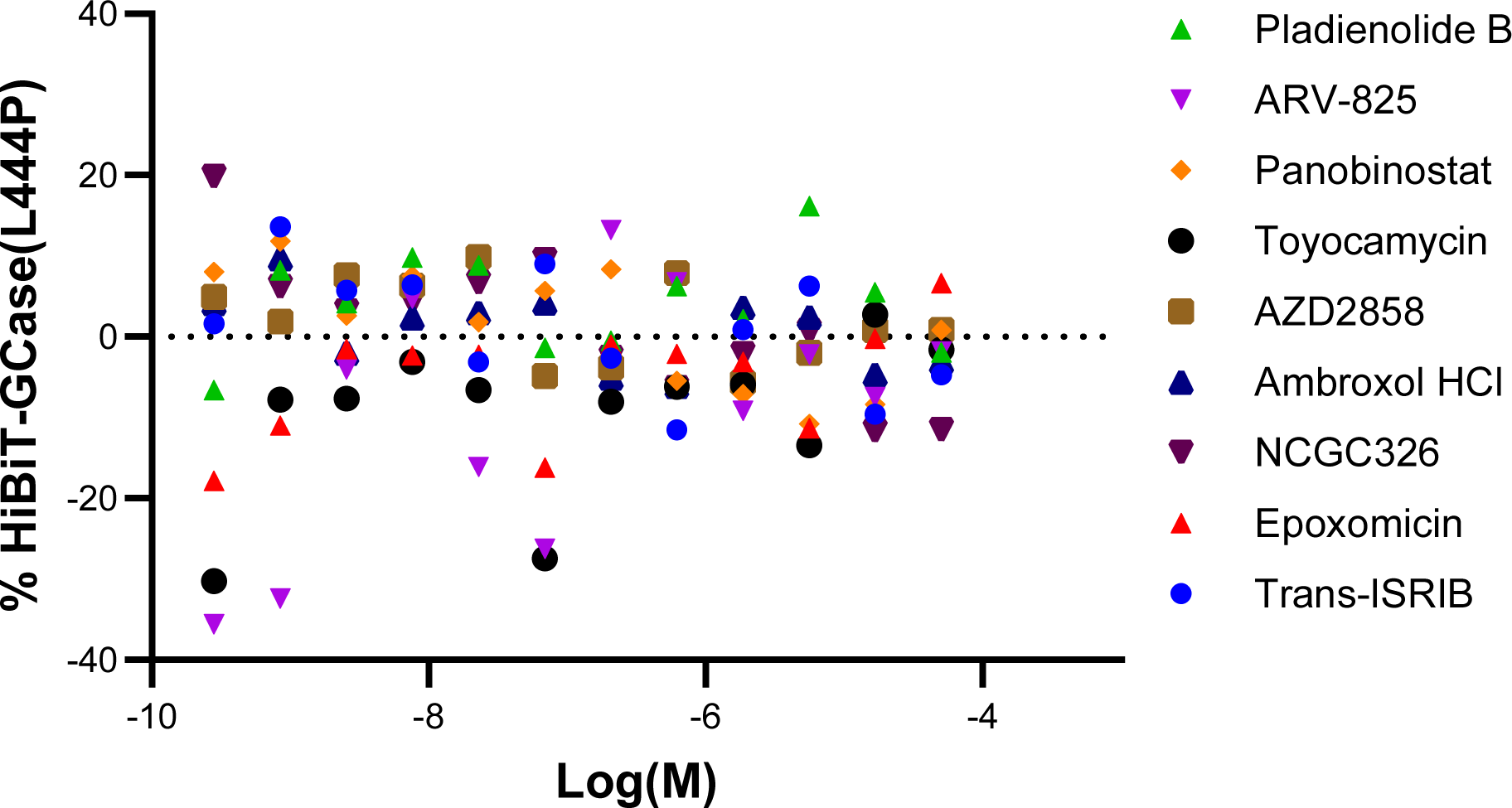
HiBiT-GCase-L444P assay interference testing for selected hits. Selected final hit compounds that increased HiBiT-GCase-L444P levels in H4 cells in the follow-up screen were tested for their ability to interfere with the reconstituted luciferase enzyme in the HiBiT-GCase assay. HiBiT-GCase-L444P H4 cells were seeded into 1536-well solid white plates (2,000 cells in 5 μL media) for 48 h and treated with a titration of compounds for 30 min, after which the HiBiT assay was performed to determine if the compounds were directly affecting HiBiT luminescence. Data are from three independent replicates.

**Figure S6.**
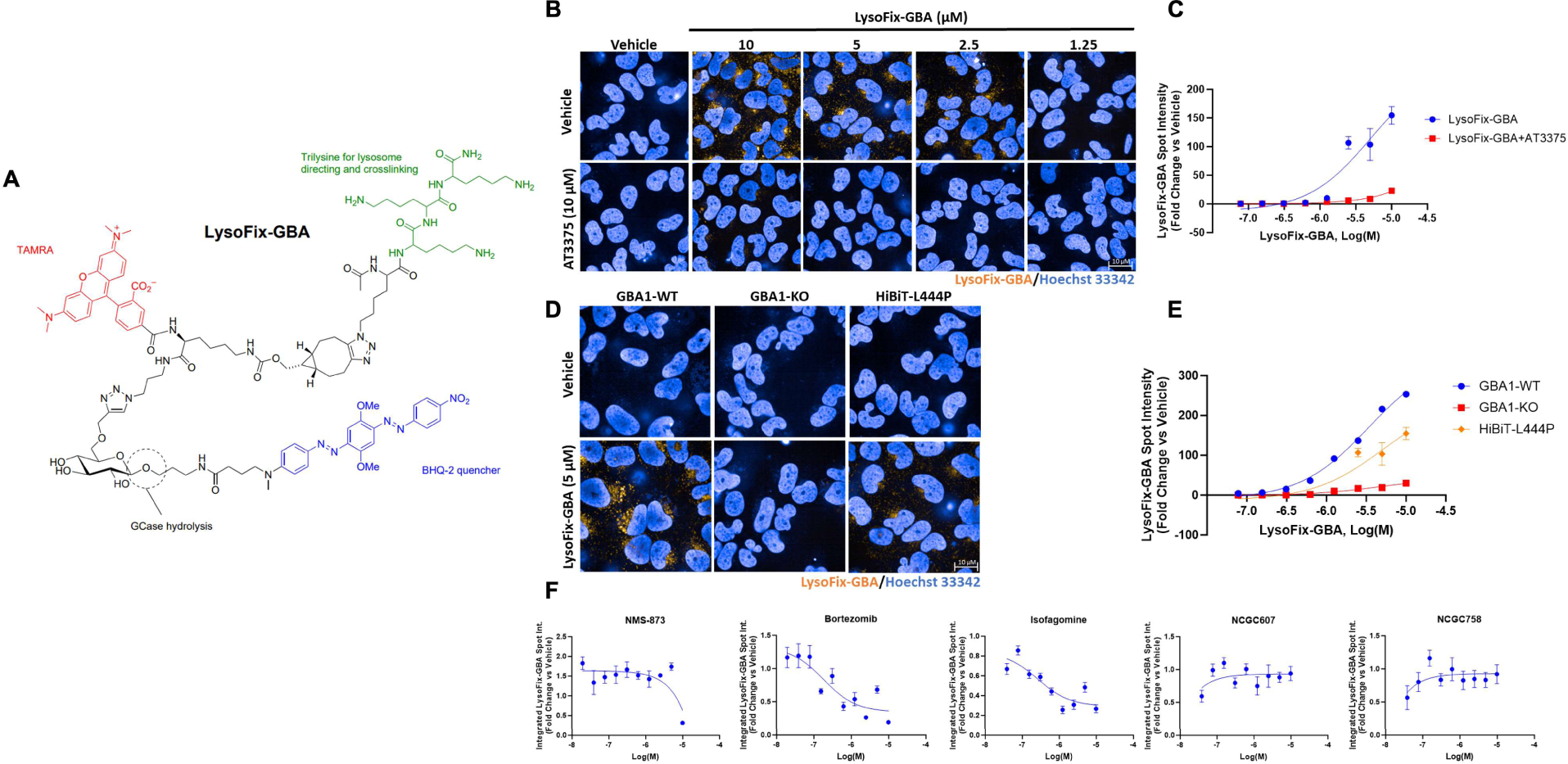
Implementation of LysoFix-GBA high-content screening assay in HiBiT-GCase-L444P H4 reporter line. (**A**) Chemical structure of LysoFix-GBA. (**B**) To optimize LysoFix-GBA concentration, HiBiT-GCase-L444P H4 cells were seeded into 384-well PerkinElmer PhenoPlates (25,000 cells in 40 μL media) for 24 h, followed by treatment with *GBA1* inhibitor AT3375 (10 μM) or vehicle (DMSO) for 24 h. Cells were then incubated with LysoFix-GBA (78 nM – 10 μM; 8-point, 2x dilution series) for 2 h at 37°C and imaged after 15 min of nuclear staining with Hoechst-33342 (1 μg/mL) in Fluorobrite media. (**C**) Data are represented as fold change in integrated LysoFix-GBA spot intensity per cell, relative to DMSO control. (Error bars: SEM [*n* = 4 – 6]). (**D, E**) Following the same approach, *GBA1*-WT, *GBA1*-KO, and HiBiT-GCase-L444P H4 cells were tested against a titration of LysoFix-GBA. (Error bars: SEM [*n* = 3]). (**F**) HiBiT-GCase-L444P H4 cells were treated with a titration of NMS-873, bortezomib, isofagomine, chaperone NCGC607, or chaperone NCGC758 for 24 h, and *GBA1* activity was assessed by LysoFix-GBA (5 μM). (Error bars: SEM [*n* = 4 – 6]).

**Figure S7.**
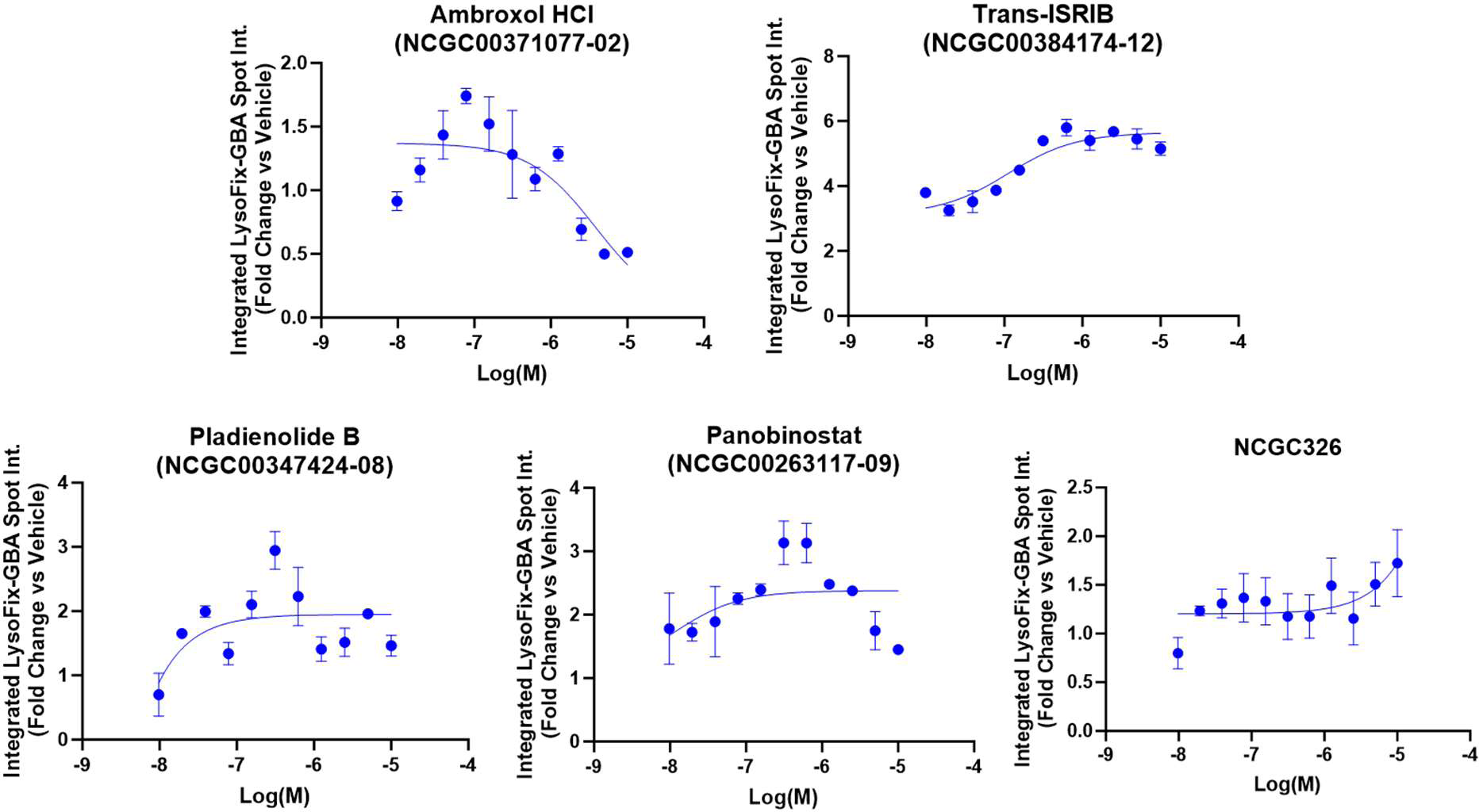
Evaluation of hit compounds with LysoFix-GBA assay after 72 h of compound incubation. HiBiT-GCase-L444P H4 cells were seeded into 384-well PerkinElmer PhenoPlates (25,000 cells in 40 μL media) and incubated for 24 h. Thereafter, the cells were treated with a titration of compounds (9.8 nM – 10 μM; 11-point, 2x dilution series) for 72 h and then incubated with LysoFix-GBA (5 μM) for 2 h at 37°C and 5% CO_2_. High-content imaging was performed after 15 min of nuclear staining with Hoechst-33342 (1 μg/mL) in Fluorobrite media. Data are represented as the fold change (compound-treated vs. DMSO-treated) in integrated LysoFix-GBA spot intensity per cell. Dose-response curves were fit using log(agonist) vs. response (three parameters). (Error bars: SEM [*n* = 2 – 5]).

**Figure S8.**
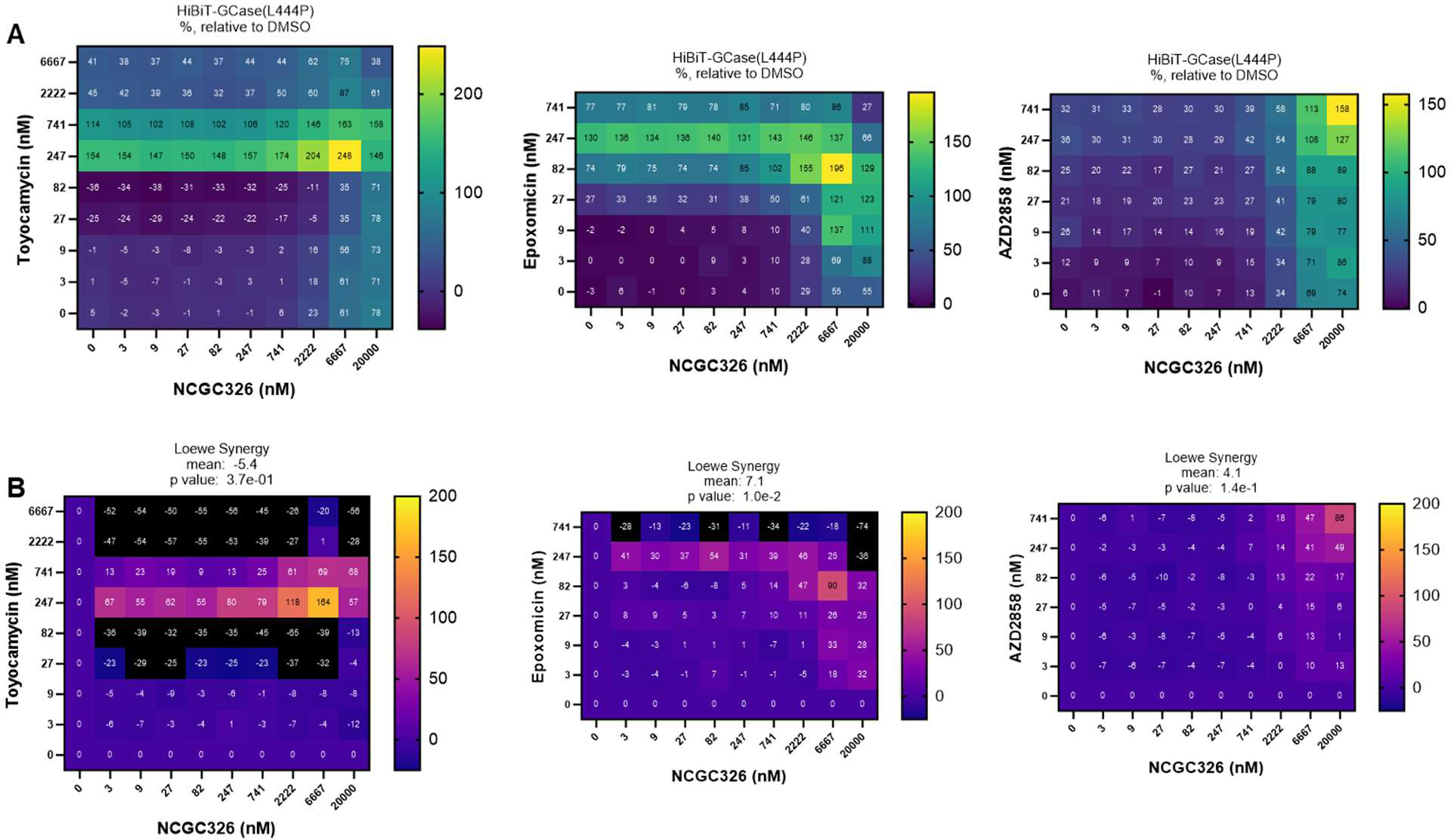
Matrix combination screening with NCGC326 and additional hits from primary HiBiT screen. HiBiT-GCase-L444P H4 cells were tested in 10 x 10 pairwise dose-response combinatorial matrix format. Cells were treated for 24 h with chaperone NCGC326 in a 9-point titration (3 nM – 20 μM, 3x dilution) against the same 9-point titration of toyocamycin, epoxomicin, or AZD2868; the HiBiT-GCase lytic assay was then performed. Luminescence response values were normalized to intraplate DMSO-treated controls, such that 100% activity reflects a doubling of HiBiT-GCase levels. Synergy was evaluated based on the dose-response matrix (**A**) and the Loewe synergy score (**B**). In general, negative, zero, and positive synergy scores indicate antagonistic, additive, and synergistic interactions, respectively, between drugs. If a dose-response curve could not be fit due to toxicity at top concentrations, these concentrations were omitted from the analysis. *n* = 3.

